# Divergent opioid-mediated suppression of inhibition between hippocampus and neocortex across species and development

**DOI:** 10.1101/2024.01.20.576455

**Authors:** Adam P. Caccavano, Anna Vlachos, Nadiya McLean, Sarah Kimmel, June Hoan Kim, Geoffrey Vargish, Vivek Mahadevan, Lauren Hewitt, Anthony M. Rossi, Ilona Spineux, Sherry Jingjing Wu, Elisabetta Furlanis, Min Dai, Brenda Leyva Garcia, Yating Wang, Ramesh Chittajallu, Edra London, Xiaoqing Yuan, Steven Hunt, Daniel Abebe, Mark A. G. Eldridge, Alex C. Cummins, Brendan E. Hines, Anya Plotnikova, Arya Mohanty, Bruno B. Averbeck, Kareem Zaghloul, Jordane Dimidschstein, Gord Fishell, Kenneth A. Pelkey, Chris J. McBain

**Affiliations:** Eunice Kennedy Shriver National Institute of Child Health and Human Development (NICHD), Section on Cellular and Synaptic Physiology, National Institutes of Health (NIH), Bethesda, MD 20892, USA; Harvard Medical School, Blavatnik Institute, Department of Neurobiology, Boston, MA 02115, USA; Stanley Center for Psychiatric Research, Broad Institute of MIT and Harvard, Cambridge, MA 02142, USA; National Institute of Mental Health (NIMH), NIH, Bethesda, MD 20892, USA; National Institute of Neurological Disorders and Stroke (NINDS) Intramural Research Program, NIH, Bethesda, MD 20892, USA

## Abstract

Within the adult rodent hippocampus, opioids suppress inhibitory parvalbumin-expressing interneurons (PV-INs), thus disinhibiting local micro-circuits. However, it is unknown if this disinhibitory motif is conserved in other cortical regions, species, or across development. We observed that PV-IN mediated inhibition is robustly suppressed by opioids in hippocampus proper but not primary neocortex in mice and nonhuman primates, with spontaneous inhibitory tone in resected human tissue also following a consistent dichotomy. This hippocampal disinhibitory motif was established in early development when PV-INs and opioids were found to regulate early population activity. Acute opioid-mediated modulation was partially occluded with morphine pretreatment, with implications for the effects of opioids on hippocampal network activity important for learning and memory. Together, these findings demonstrate that PV-INs exhibit a divergence in opioid sensitivity across brain regions that is remarkably conserved across evolution and highlights the underappreciated role of opioids acting through immature PV-INs in shaping hippocampal development.

## INTRODUCTION

The endogenous opioid system in the CNS plays a crucial role in pain sensation, stress response, and mood^1,2^ and is a key target for exogenous drugs of abuse including heroin and fentanyl. With a high risk of dependency, mortality from respiratory depression, and epidemic levels of abuse, opioids have fueled a public health emergency. One third of pregnant women report opioid use, and in almost 60,000 annual pregnancies there is reported opioid abuse^3^. Children born to opioid-dependent mothers are at increased risk of neurodevelopmental deficits within cognitive, psychomotor, and language domains^4^. Despite these risks, there remains an urgent need for potent and safe analgesics. To develop improved opioid or non-opioid analgesics, research is required into the mechanisms of opioids at the cellular and microcircuit level. Novel genetic tools enable us to study the mechanisms of opioids in distinct neuronal subpopulations across species, with the goals of elucidating endogenous opioid function and developing more targeted treatments.

Endogenous opioids (e.g., endorphins, enkephalins) and exogenous opiates (e.g., morphine, fentanyl) act with differing specificities on mu, delta, and kappa opioid receptors (μOR, δOR, and κOR). Notably, μORs are associated with reward-processing of both natural stimuli and drugs of abuse^1,5,6^. These G_i_/G_o_ coupled metabotropic receptors have both pre- and post-synaptic inhibitory secondary effects^7,8^ including the pre-synaptic closure of Ca^2+^ channels^9,10^ and post-synaptic opening of inward-rectifying K^+^ channels^11–13^. Within the hippocampus (HPC), μOR activation has an overall disinhibitory effect on the network^14–17^ due to a preferential suppression of inhibitory GABAergic cells^18–20^. Within the CA1 region of HPC, this suppression of inhibition has been primarily associated with parvalbumin-expressing interneurons (PV-INs)^21–23^. Perisomatic-targeting PV-INs of the HPC highly express μORs/δORs, though not exclusively, as lower percentages of dendritic-targeting somatostatin-expressing interneurons (SST-INs) and neuropeptide Y-expressing ivy and neurogliaform cells (NPY-INs) also express μORs/δORs^24,25^, and are suppressed by opioids^26,27^. Although μORs/δORs are expressed throughout the CNS^28–32^, most functional studies of opioid suppression of interneurons have been restricted to the adult rodent HPC, with less focus on other regions, species, or across development.

PV-INs are critical organizers of rhythmic activity important for learning and memory including sharp wave ripples (SWRs) and gamma oscillations^33^. PV-INs throughout the forebrain have common developmental origins in the medial ganglionic eminence (MGE)^34^ and are often treated as monolithic with common circuit motif functionality, leading to textbook observations of inhibitory microcircuits that may not generalize across brain regions^33,35,36^. PV-INs within different brain regions may express unique receptors, as is the case in striatum, where PV-INs express the CB_1_ cannabinoid receptor^37^, which in HPC is selectively expressed by cholecystokinin-expressing interneurons (CCK-INs)^38^. Given the degree of inhibitory control this neuronal population has over cortical microcircuits, any regional specializations become essential to understand if PV-INs are expected to participate in or be targets of therapeutic interventions for addiction. Moreover, it is essential to translate any rodent findings to higher species as cellular and circuit motifs may have diverged over 70 million years of evolution, particularly considering evidence of human-specific PV-IN innovations in channel expression and electrophysiological properties^39,40^.

In the present study we examined the opioid-mediated suppression of PV-INs across cortical regions, species, and development. We observed this disinhibitory motif was unique to the hippocampus proper (CA1-3), was remarkably conserved from mouse to macaque to human, and was established in early postnatal development, just as inhibitory synapses were being established, with important control over population activity of the developing hippocampus. The hippocampal specificity of opioid-mediated disinhibition has profound implications for rhythmic activities supporting learning and memory. Indeed, prior studies have demonstrated that both SWRs^41^ and gamma oscillations^42^ are highly sensitive to opioid administration. In the present study we extend these findings to opioid modulation of postnatal population activity, with severe implications for the harmful aspects of opioid use *in utero* and early development. Together, our findings demonstrate that despite common developmental origins in the MGE, not all PV-INs are destined to fulfill equivalent circuit roles.

## RESULTS

### μORs are selectively enriched in hippocampal PV-INs

μORs are expressed throughout hippocampus (HPC) and neocortex (CTX)^1^, and within the HPC are highly enriched in PV-INs^24,25^. However, less is known about the specificity of μOR expression to PV-INs throughout CTX. To address this, we compared the expression of *Oprm1* (encoding μORs) across HPC and CTX in GABAergic INs via single-nucleus RNA sequencing (snRNAseq) of postnatal day (P)28 mice, observing an enrichment of *Oprm1* in HPC versus CTX within the delineated PV cell cluster (Fig. 1A-B). We next assessed the spatial colocalization of mRNA via RNAscope, observing a decreased percentage of *Pvalb*+ somata colocalized with *Oprm1* in neocortical pyramidal cell layers (both supragranular layers 2/3 and infragranular layers 5/6) in primary motor (M1), somatosensory (S1) and visual (V1) cortex relative to hippocampal pyramidal cell layers in CA (CA1-3, Fig. 1C-D). Notably, we observed several *Oprm1*+*Pvalb*-cells in CTX (Fig. 1C, *open arrow*), which we did not encounter in HPC. To assess if transcript associated with the translational machinery was altered, we employed a RiboTag sequencing approach of all MGE-derived interneurons (MGE-INs, of which PV-INs comprise a substantial fraction^43^), employing Nkx2.1^Cre/+^:Rpl22(RiboTag)^HA/HA^ mice^44–46^. We observed a selective enrichment of *Oprm1* in hippocampal MGE-INs relative to neocortical MGE-INs and all bulk hippocampal/neocortical tissue (Fig. 1E).

**Fig. 1:**
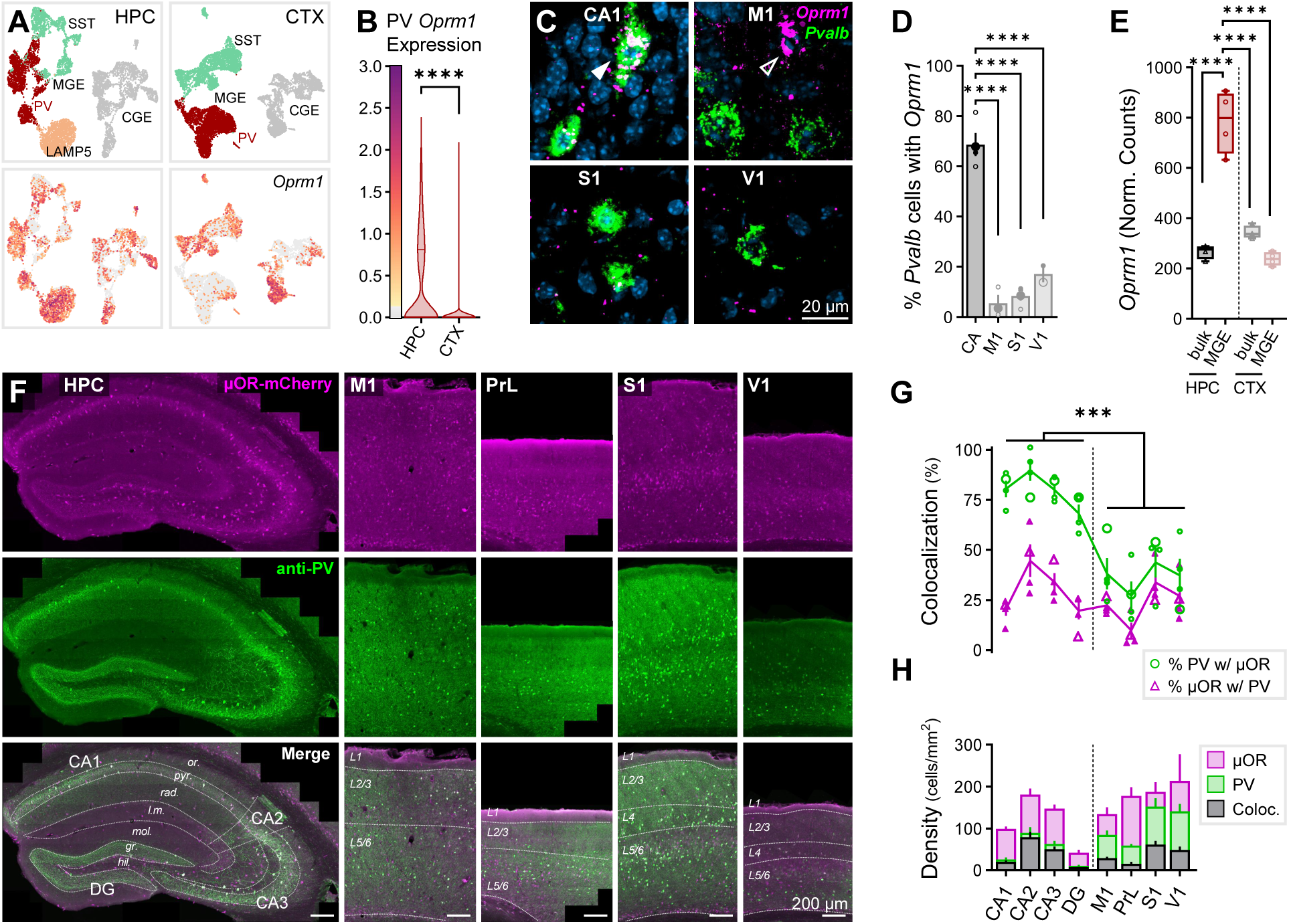
μORs are selectively enriched in hippocampal PV-INs. (**A**) snRNAseq of GABAergic INs in P28 mice across HPC and CTX, highlighting *Oprm1* expression. Cardinal clusters of Lhx6-expressing MGE-INs are colored and delineated as SST (*aquamarine*), PV (*red*), and LAMP5 (*peach*) in contrast to caudal ganglionic eminence (CGE)-derived INs (gray). (**B**) *Oprm1* expression of PV cluster cells, n_cell_ = 1920 (HPC), 3935 (CTX) from n_mice_ = 2 (1F), age = P28. Asterisks represent Dunn’s *post hoc* comparisons of all HPC/CTX differences for *Oprm1*/*Oprd1* in PV/SST-INs (see Fig. 4B for other comparisons). (**C**) *In situ* hybridization (ISH) RNAscope for *Oprm1* and *Pvalb* in wild-type (WT) mice across pyramidal cell layers in CA (CA1-3) regions of HPC and primary neocortical regions M1, S1, V1. Closed arrow (CA1) indicates *Oprm1*+*Pvalb*+ colocalized cell, while open arrow (M1) indicates *Oprm1*+*Pvalb*-cell. (**D**) Colocalization quantification of percentage of *Pvalb*+ somata co-expressing *Oprm1* for 2-5 sections from each of n = 4 (2F) WT mice, age = P62. Asterisks represent Tukey’s *post hoc* comparisons after a significant effect of region was observed via 1-way ANOVA. (**E**) RiboTag-associated *Oprm1* expression in n = 4 (2F) P120 Nkx2.1^Cre^: Rpl22(RiboTag)^HA/HA^ mice, comparing bulk HPC/CTX tissue to MGE-INs. Asterisks represent Tukey’s *post hoc* comparisons after significant effects of region, cell type, and region × cell type interaction were observed via 2-way ANOVA. (**F**) Immunohistochemical (IHC) stain for PV in μOR-mCherry mice (boosted with anti-RFP) labeling hippocampal regions: CA1, CA2, CA3, and DG and layers: *stratum* (*str*.) *oriens* (*or*.), *pyramidale* (*pyr*.), *radiatum* (*rad*.), *lacunosum-moleculare* (*l.m.*), *molecular* (*mol*.), *granular* (*gr*.), *hilus* (*hil*.), and neocortical regions: M1, PrL, S1, V1, and layers: L1, L2/3, L4, L5/6. (**G-H**) Quantification of IHC for n_section_ = 4 from n_mice_ = 2F, age = P70 for all layers of indicated region. (**G**) Colocalization quantification of the percent of PV cells co-expressing μORs (*green circles*) and percent of μOR cells co-expressing PV (*magenta triangles*). Enlarged markers indicate quantification from example images. Asterisks represent significance of unpaired t-test comparing the percentage of PV cells co-expressing μORs across all HPC and CTX. (**H**) Cell density (cells/mm^2^) for μOR-expressing cells (*magenta*), PV cells (*green*) and PV-μOR colocalized cells (*black*). Data are represented as mean ± SEM. * p < 0.05, ** p < 0.01, *** p < 0.001, **** p < 0.0001. See *Table S1* for statistical details for this and all subsequent figures.

At the protein level, we quantified PV-μOR colocalization via immunohistochemistry (IHC) by staining for PV in P70 μOR-mCherry mice (Fig. 1F). μOR-mCherry mice were employed as their expression pattern closely matched μOR antibody signal with brighter somatic labeling (Fig. S1A-B). We observed an increased colocalization of PV and μOR somata across all layers of hippocampal proper (CA1, CA2, CA3) relative to primary cortical regions (M1, S1, V1) and higher-order prelimbic cortex (PrL) (Fig. 1G-H). Hippocampal dentate gyrus (DG) exhibited intermediate PV-μOR colocalization relative to hippocampus proper and neocortex.

As hippocampal MGE-INs are born earlier than neocortical MGE-INs (embryonic day (E)11.5^43^ versus E13.5^47,48^), we next examined if embryonic PV-IN birthdate was associated with *Oprm1* expression. Through EdU proliferation labeling across E11-15 (Fig. S2A-C), we replicated prior observations that hippocampal versus neocortical PV-INs are born earlier (Fig. S2D). This large biological replicate RNAscope study also fully supported the above RNAscope evaluation of weak *Oprm1* expression within cortical versus hippocampal PV-INs. However, later-born PV-INs were no more likely to express *Oprm1* (Fig. S2E), suggesting factors other than embryonic birthdate establish the hippocampal-neocortical divergence in PV-IN μOR expression.

### Hippocampal PV-INs are selectively hyperpolarized by opioids

μORs hyperpolarize neurons and reduce neurotransmitter release via combined interactions with inward-rectifying K^+^ channels and closure of presynaptic Ca^2+^ channels^7,8^. To determine whether a functional divergence existed between hippocampal and neocortical PV-INs, we recorded in a whole-cell configuration, currents elicited by the administration of μOR agonist DAMGO (100 nM) and antagonist CTAP (500 nM) in PV-tdTomato^+/-^ mice (Fig. 2A-B). Consistent with prior studies^21,22^, we observed an outward DAMGO-mediated hyperpolarizing current in CA1 PV-INs voltage-clamped to -50 mV (ΔV = +60 ± 12 pA, Fig. 2C, D_1_-F_1_). However, no significant change in holding current was observed in V1 PV-INs (+12 ± 5 pA, Fig. 2C, D_2_-F_2_). Of the CA1 PV-INs with sufficiently recovered morphology to assess axonal target (n_cell_ = 5 of 11), 3 exhibited perisomatic-targeting basket cell (BC) morphology, 1 bistratified (BSC) morphology targeting apical and basal dendrites, and 1 axo-axonic (AAC) morphology targeting the axon-initial segment (AIS) of PCs. The hyperpolarizing DAMGO-elicited current was observed in all PV-IN subpopulations (BC: +73 ± 37, BSC: +85, AAC: +101 pA), which is of interest as prior research has suggested AACs are less sensitive than BCs to DAMGO following carbachol administration^42^. V1 PV-INs had somata spanning cortical layers 2 through 6, and of those with sufficiently labeled axons (n_cell_ = 6/15), 5 exhibited BC and 1 AAC morphology, with neither subpopulation responsive to DAMGO. The input resistance was also decreased in CA1 PV-INs (Fig. 2G_1_), consistent with the opening of ion channels, which was not observed in V1 PV-INs (Fig. 2G_2_). We next substituted GTP in the internal pipette solution with the non-hydrolyzable GTP analogue GTPγS, which binds to and prevents further GPCR activation^49^, and observed no DAMGO-mediated change in holding current in either CA1 or V1 (Fig. 2H-J), indicating a direct mechanism of GPCRs expressed in the recorded PV-INs. Together, these findings point to a transcriptional, translational, and functional enrichment of μORs within hippocampal versus neocortical PV-INs.

**Fig. 2:**
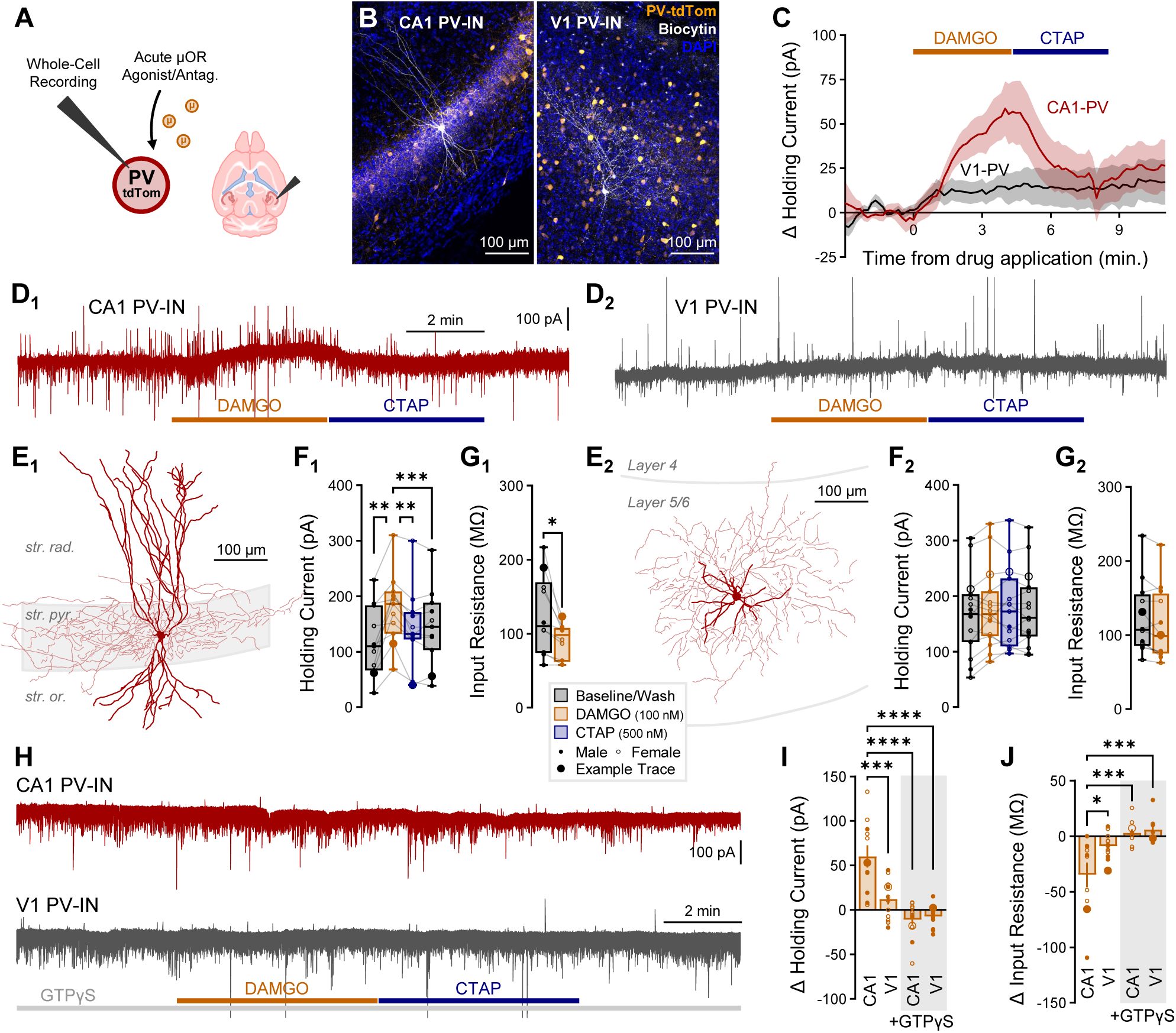
Hippocampal PV-INs are selectively hyperpolarized by opioids. (**A**) Schematic of whole-cell recordings of PV-INs from PV-tdTom mice voltage-clamped to -50 mV to record the effect of μOR agonist/antagonist DAMGO/CTAP. (**B**) Representative *post hoc* staining of recorded PV-INs in CA1 and V1. (**C**) Average change in holding current elicited by DAMGO/CTAP normalized to baseline for n_cell_ = 11 CA1 from n_mice_ = 5 (3F), age = P37-70 (P57 ± 7) and n_cell_ = 15 V1 from n_mice_ = 4 (3F), age = P47-71 (P58 ± 6), with (**D**) example traces showing effects of drugs on baseline holding current (rapid inward and outward currents reflect sEPSCs/sIPSCs), (**E**) *post hoc* reconstructions, (**F**) holding current summary data, and (**G**) input resistance summary data in a subset of cells, for CA1 PV-INs (**1***, left*) and V1 PV-INs (**2***, right*). Asterisks represent Tukey’s *post hoc* comparisons after a significant effect of treatment was found via 1-way repeated measures ANOVA. (**H**) Example traces showing effect of DAMGO/CTAP with GTPγS substituted for GTP in the internal pipette solution, with DAMGO administration occurring 15 min. after break-in, performed in n_cell_ = 12 CA1 and 12 V1 from n_mice_ = 4 (2F), age = P118-392 (P315 ± 66). Summary statistics for DAMGO-mediated change from baseline in (**I**) holding current and (**J**) input resistance for the four conditions (CA1/V1 and GTP/GTPγS). Asterisks represent Tukey’s *post hoc* comparisons after significant effects of region, internal solution, and region × internal interaction were observed via 2-way ANOVA. In these and all subsequent plots, data from male/female subjects are indicated with a closed/open marker, with enlarged marker indicating example trace.

### Opioids selectively suppress hippocampal proper but not neocortical PV-IN mediated inhibition

To determine if the selective enrichment of μORs within hippocampal PV-INs reduces inhibitory synaptic transmission, we adopted an optogenetic approach to light-activate PV-INs and record their output in downstream pyramidal cells (PCs) in PV^Cre/+^:ChR2^fl/+^ mice (Fig. 3A-B). GABA_A_-mediated currents were pharmacologically isolated with bath administration of AMPA, NMDA, and GABA_B_ antagonists (µM: 10 DNQX, 50 DL-APV, and 1 CGP55845 respectively), with drugs of interest (nM: 100 DAMGO, 500 CTAP) bath administered for at least 5 minutes. Consistent with prior hippocampal studies^21,22^, light-evoked inhibitory post-synaptic currents (leIPSCs) were suppressed by DAMGO in CA1-PCs (Fig. 3C). To assess a pre- or post-synaptic mechanism, we analyzed the coefficient of variation (Fig. 3C), and in a subset of cells, the paired pulse ratio (Fig. 3D), observing an increase in both metrics (Fig. 3E-F), consistent with a presynaptic mechanism and prior studies^21,23^. Notably however, PV-IN output was differentially modulated between HPC and CTX, with significant suppression (64 ± 5% of baseline) observed in CA1-PCs and no suppression (108 ± 6%) observed in M1-PCs (Fig. 3G). To broadly assess this across cortical regions, we recorded leIPSCs in three hippocampal (CA1, CA3, DG) and four neocortical regions (M1, PrL, S1, V1). Additionally, as differences in PV connectivity have been reported between *radiatum* (*rad.*)-adjacent superficial and *oriens* (*or.*)-adjacent deep PCs (sPCs and dPCs respectively)^50,51^, we segregated recordings from these cell populations. DAMGO suppressed leIPSCs in both CA1-sPCs (55 ± 8%, Fig. 3H) and CA1-dPCs (72 ± 7%, Fig. 3I), with no difference between these cell populations (p = 0.135, Fig. S3A). Both in CA1-sPC and CA1-dPC recordings, CTAP reversed the DAMGO-mediated suppression, although this could be achieved through wash alone (data not shown). DAMGO also suppressed leIPSCs in CA3-PCs (69 ± 6%), although this did not fully reverse with either CTAP or wash (Fig. 3J). Interestingly, this long-lasting μOR-mediated suppression in CA3 is similar to observations of a long-lasting δOR-mediated suppression in CA2 that was not observed in CA1^52^. In DG granule cells (DG-GCs), we observed a highly variable response, with no significant group effect of DAMGO or CTAP (95 ± 8%, Fig. 3K). Throughout neocortex, we also observed highly variable responses with no significant group effects of either DAMGO or CTAP in M1 (108 ± 6%, Fig. 3L), PrL (96 ± 8%, Fig. 3M), S1 (107 ± 9%, Fig. 3N), and V1 (89 ± 11%, Fig. 3O). Neocortical PCs were recorded in both supragranular (2/3) and infragranular (5/6) layers. To address the possibility that differing opioid sensitivity between cortical layers contributes to the increased variability in the observed leIPSC amplitude, we combined primary cortex (M1, S1, V1) PCs and segregated the combined data across L2/3 and L5/6, with no differences observed in the baseline normalized DAMGO response (p = 0.291, Fig. S3B). We also explored the increased variability of cortical responses by removing outliers (ROUT method) and all cells with greater than 150% increase from baseline, but still observed no significant group effects (data not shown).

**Fig. 3:**
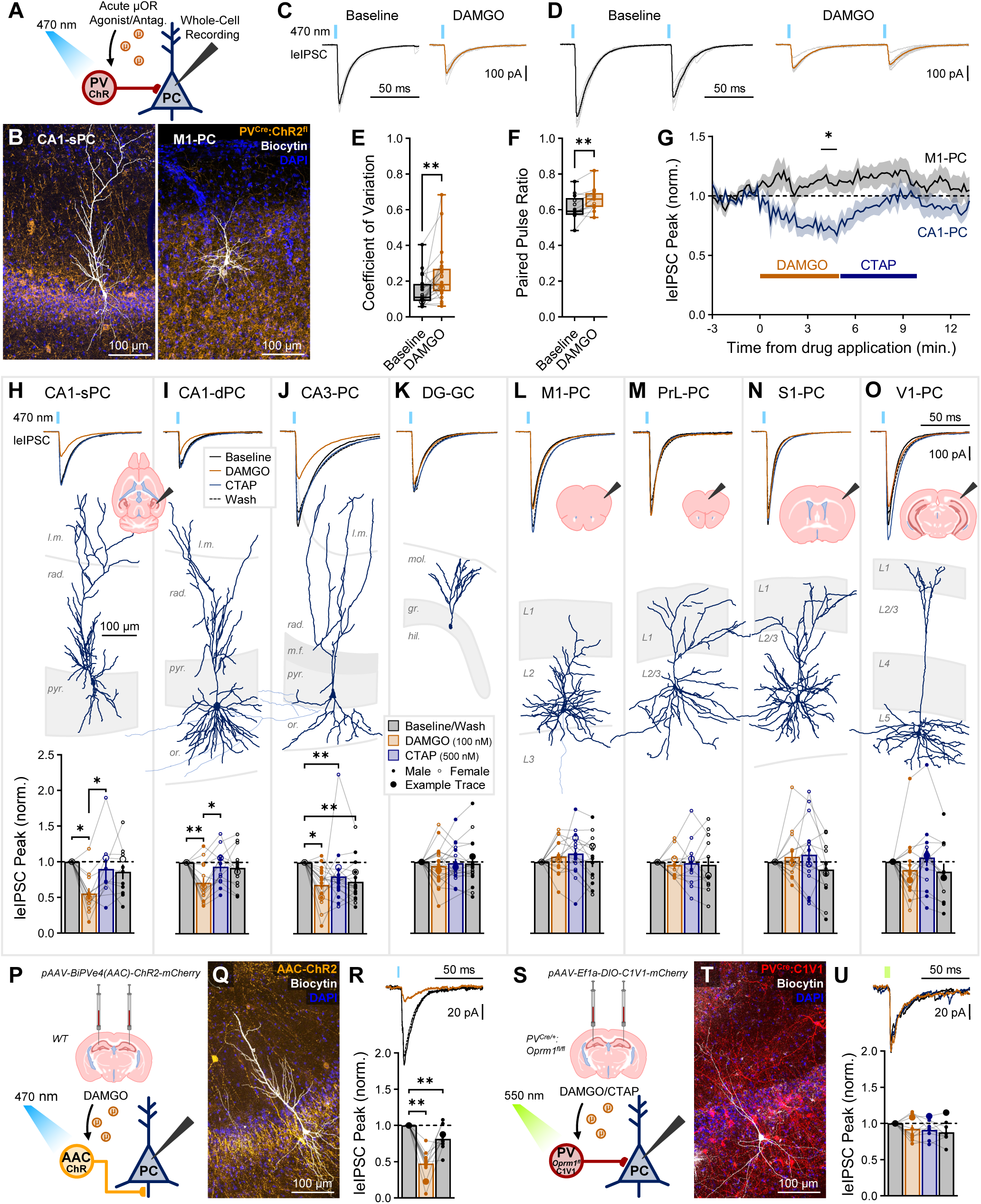
Opioids selectively suppress hippocampal but not neocortical PV-IN inhibition. (**A**) Schematic of whole-cell recordings of PCs from PV^Cre/+^:ChR2^fl/+^ mice voltage-clamped at -70 mV (with high Cl^-^ internal) to record the effect of DAMGO/CTAP on light-evoked IPSCs (leIPSCs). (**B**) Representative *post hoc* staining of a recorded CA1 superficial PC (sPC) and M1 layer 2/3 PC. (**C**) Example CA1-PC leIPSC traces, dark lines represent average of ten light grey traces, showing an increased coefficient of variation (CV = SD/mean). (**D**) Example paired pulse ratio (PPR = peak2/peak1) traces recorded in a subset of CA1-PCs. Summary data for (**E**) CV for n_cell_ = 24 from n_mice_ = 10 (5F), age = P28-133 (P55 ± 9) and (**F**) PPR for n_cell_ = 13 from n_mice_ = 3 (2F), age = P78-83 (P80 ± 2). Asterisks represent results of paired t-test/Wilcoxon. (**G**) Average leIPSC peak amplitude normalized to baseline in response to DAMGO/CTAP for n_cell_ = 19 CA1-PCs from n_mice_ = 7 (4F), age = P28-55 (P44 ± 3) and n_cell_ = 16 M1-PCs from n_mice_ = 6 (4F), age = P34-61 (P46 ± 5). Asterisk marks regions where data (10 s bins) survived multiple comparisons via 2-way ANOVA. (**H**-**O**) Mouse leIPSC experiments with (*top*) averaged example traces, (*top inset*) example of brain slices, (*middle*) representative *post hoc* reconstructions, and (*bottom*) summary data in (**H**) n_cell_ = 12 CA1-sPCs from n_mice_ = 6 (3F), age = P28-133 (P62 ± 15), (**I**) n_cell_ = 14 CA1 deep PCs (dPCs) from n_mice_ = 9 (5F), age = P28-133 (P54 ± 10), (**J**) n_cell_ = 17 CA3-PCs from n_mice_ = 7 (2F), age = P41-80 (P65 ± 6), (**K**) n_cell_ = 21 DG granule cells (GCs) from n_mice_ = 9 (3F), age = P41-125 (P69 ± 9), (**L**) n_cell_ = 16 M1-PCs from n_mice_ = 6 (4F), age = P34-61 (P46 ± 5), (**M**) n_cell_ = 12 PrL-PCs from n_mice_ = 3 (2F), age = P34-144 (P107 ± 37), (**N**) n_cell_ = 15 S1-PCs from n_mice_ = 4 (3F), age = P45-52 (P49 ± 2), and (**O**) n_cell_ = 16 V1-PCs from n_mice_ = 6 (2F), age = P27-133 (P73 ± 18). (**P**) Viral bilateral intra-cerebral injection of *pAAV-BiPVe4(AAC)-ChR2-mCherry* in dorsal hippocampus of WT mice, and schematic of leIPSC recording. (**Q**) Representative *post hoc* staining of a recorded CA1-PC (*white*) and viral expression (*orange*, see *Fig. S4* for AAC viral validation). (**R**) Summary data for n_cell_ = 12 CA1-PCs from 1M, age = P50. (**S**) Viral bilateral injection of Cre-dependent *pAAV-Ef1a-DIO-C1V1-mCherry* in dorsal hippocampus of *PV^Cre/+^:Oprm1^fl/fl^* mice, and schematic of leIPSC recording. (**T**) Representative *post hoc* staining of a recorded CA1-PC (*white*) and viral expression (*red*). (**U**) Summary data for n_cell_ = 9 CA1-PCs from 2M, age = P58-61. Asterisks in (*H-U*) summary plots represent *post hoc* comparisons (Tukey’s/Dunn’s) after a significant effect of treatment was found via 1-way repeated measure ANOVA/mixed model/Friedman’s test.

We next examined the role of sex, as there have been reported differences in morphine response^53^ and µOR trafficking in hippocampal PV-INs^54,55^. Most regions were insufficiently powered to segregate along sex. However, by combining hippocampal proper (CA1, CA3) and primary neocortex (M1, S1, V1), we achieved sufficiently powered groups to analyze via 2-way ANOVA the effects of sex and region. A significant effect of region was found with sex not reaching significance (Fig. S3C). Thus, sex did not appear to be a driving confounder behind the observed hippocampal-neocortical differences.

The concentration of DAMGO was selected to be consistent with prior literature^21^, but to determine if the lack of cortical effect was due to lower affinity in these regions, we increased the DAMGO concentration 10-fold (1 μM) and observed similar results, with suppression in CA1 (58 ± 8% of baseline) and none in DG (96 ± 11%) or M1 (108 ± 9%, Fig. S3D). These findings were not unique to the selective μOR agonist DAMGO, as the less selective and potent μOR agonist morphine (10 μM) had similar effects in CA1 (56 ± 10%) and S1 (110 ± 12%, Fig. S3E). To test if endogenous opioids were tonically suppressing PV-IN release, we applied neutral antagonist CTAP preceding DAMGO, observing no increase from baseline in CA1 (83 ± 10%) or S1 (99 ± 13%), nor any effect from subsequent DAMGO administration with CTAP still applied (Fig. S3F). As μORs and δORs can become constitutively active, in which they activate G proteins even in the absence of agonist^56,57^, we also tested the antagonist naloxone, which can function as an inverse agonist after opioid exposure^58,59^. We observed no increase from baseline in PV-IN output from 1 μM naloxone in CA1 (110 ± 18%) or S1 (81 ± 11%, Fig. S3G). All drug conditions are summarized in Fig. S3H.

Although we previously observed DAMGO-mediated hyperpolarization in one morphologically identified PV-AAC, we more thoroughly explored this cell population due to prior reports of divergent opioid response amongst BCs and AACs^42^. We employed the novel BiPVe4(AAC) enhancer virus (see Fig. S4 and^60^ for validation), which labels AACs with closely matched features as other strategies to label this subpopulation{Dudok, 2021 #1651;Raudales, 2023 #1973. After injecting WT mice with *pAAV-BiPVe4(AAC)-ChR2-mCherry*, we observed a robust DAMGO-mediated suppression of AAC inhibition in CA1 (47 ± 5%, Fig. 3P-R). Moreover, we confirmed DAMGO-mediated outward currents in a larger cohort of AACs (Fig. S4F-H). Critically, suppression of PV-IN inhibition was directly mediated by μORs expressed within PV-INs, as we observed no DAMGO-mediated suppression (92 ± 6%) following selective knockout of *Oprm1* in PV-INs through breeding PV^Cre/+^:*Oprm1*^fl/fl^ and probing output from these mice after injection with the red-shifted opsin C1V1 under control of Cre-recombinase (Fig. 3S-U and Fig. S5).

### δOR activation and SST-IN inhibition exhibit similar functional HPC-CTX divergence

Hippocampal PV-INs are also enriched in δORs, with μORs and δORs functioning through partially occlusive downstream pathways to hyperpolarize interneurons{He, 2021 #1680}. In contrast to our observed lack of *Oprm1* expression in neocortical PV-INs, we observed a high level of *Oprd1* transcript within PV-INs in our snRNAseq dataset (Fig. 4A-B), as well as high neocortical *Pvalb*-*Oprd1* colocalization via RNAscope, albeit significantly less than hippocampal *Pvalb*-*Oprd1* colocalization (Fig. 4C-D). Our RiboTag dataset revealed consistent findings; both hippocampal and neocortical MGE-INs were enriched in *Oprd1*, with higher enrichment in hippocampal versus neocortical MGE-INs (Fig. 4E). We next applied the δOR-selective agonist DPDPE (500 nM) in PV^Cre/+^:ChR2^fl/+^ mice (Fig. 4F), observing a significant suppression in CA1 (76 ± 5%) that did not reach significance in V1 (88 ± 6%, Fig. 4G). However, when analyzing all normalized drug responses with a less conservative 1-way rather than repeated-measures ANOVA of the raw data, we found that across all drug conditions in CTX, only DPDPE resulted in a significant suppression (Fig. S3H). Thus, δOR activation may modestly suppress neocortical PV-IN output.

**Fig. 4:**
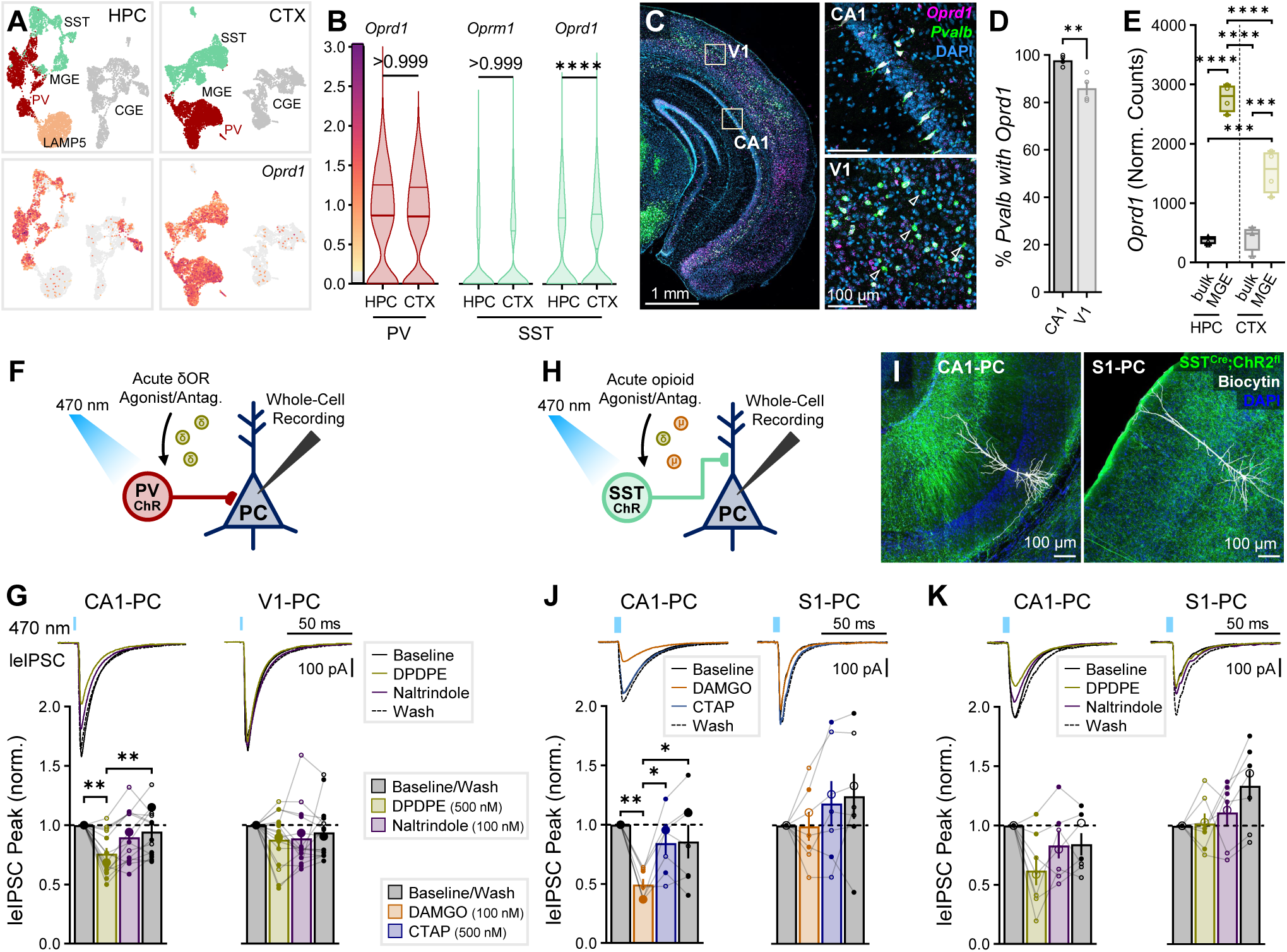
δOR activation and SST-IN inhibition exhibit similar functional HPC-CTX divergence. (**A**) snRNAseq of cardinal clusters of HPC/CTX MGE-INs, highlighting *Oprd1* expression. (**B**, *left*) *Oprd1* expression in PV cluster cells, n_cell_ = 1920 (HPC), 3935 (CTX) and (*right*) *Oprm1* and *Oprd1* expression in SST cluster cells, n_cell_ = 1518 (HPC), 3002 (CTX), from n_mice_ = 2 (1F), age = P28. Asterisks and non-significant p-values represent results of Dunn’s *post hoc* comparisons after comparing all HPC/CTX differences for *Oprm1*/*Oprd1* in PV /SST-INs (including from Fig. 1B). (**C**) RNAscope for *Oprd1* and *Pvalb* in WT mice comparing CA1 to V1 (all layers). Majority of cells were *Oprd1+Pvalb+* colocalized (not indicated), with *Oprd1-Pvalb+* cells indicated with open arrows. (**D**) Colocalization quantification of percentage of *Pvalb*+ somata co-expressing *Oprd1* for 1 section each from n = 4 (2F) WT mice, age = P30-32. Asterisks represent significance of unpaired t-test. (**E**) RiboTag-associated *Oprd1* expression in n = 4 (2F) P120 Nkx2.1^Cre^: Rpl22(RiboTag)^HA/HA^ mice, comparing bulk HPC/CTX tissue to MGE-INs. Asterisks represent Tukey’s *post hoc* comparisons after significant effects of region, cell type, and region × cell type interaction were observed via 2-way ANOVA. (**F**) Schematic of whole-cell recordings of PCs recorded in PV^Cre/+^:ChR2^fl/+^ mice voltage-clamped at -70 mV (with high Cl^-^ internal) to record the effect of δOR agonist/antagonist DPDPE/naltrindole on leIPSCs. δOR leIPSC experiments were performed in (**G**) n_cell_ = 14 CA1-PCs and 15 V1-PCs from n_mice_ = 7 (2F), age = P57-90 (P73 ± 5), with (*top*) averaged example traces and (*bottom*) summary data. (**H**) Schematic of whole-cell recordings of PCs voltage-clamped at -70 mV recorded in SST^Cre/+^:ChR2^fl/+^ mice to record the effect of µOR/δOR agonists/antagonists on leIPSCs. (**I**) Representative *post hoc* staining of recorded CA1 and S1 PCs in SST^Cre/+^:ChR2^fl/+^ mice. SST leIPSC experiments with (**J**) DAMGO/CTAP were performed in n_cell_ = 7 CA1-PCs and 7 S1-PCs from n_mice_ = 4 (3F), age = P57-240 (P162 ± 46) and with (**K**) DPDPE/Naltrindole in n_cell_ = 7 CA1-PCs and 7 S1-PCs from n_mice_ = 4 (2F), age = P74-135 (P106 ± 13), with (*top*) averaged example traces and (*bottom*) summary data. Asterisks in (*G, J, K)* represent Tukey’s/Dunn’s *post hoc* comparisons after a significant effect of treatment was found via 1-way repeated measure ANOVA/Friedman’s test.

Somatostatin-expressing interneurons (SST-INs) comprise a substantial portion of MGE-INs^43^, and within the HPC also express μORs and δORs^25^. Both in HPC and CTX, snRNAseq revealed an apparent enrichment of *Oprm1* and *Oprd1* within SST-INs (Fig. 1A, Fig. 4A,B). We undertook similar optogenetic experiments in SST^Cre/+^:ChR2^fl/+^ mice (Fig. 4H-I), observing that as with PV-INs, leIPSCs from SST-INs were reversibly suppressed by DAMGO in CA1 (49 ± 5%) but not S1 (99 ± 12%, Fig. 4J). DPDPE resulted in a non-significant suppression in CA1 SST-IN output (62 ± 12%) with no suppression in S1 (101 ± 8%, Fig. 4K). Thus, hippocampal SST-INs appear to exhibit a similar functional specialization as PV-INs regarding opioid response, despite an apparent enrichment of μORs and δORs within neocortical SST-INs. The lack of effect suggests altered subcellular localization with limited opioid receptor expression at PV/SST-IN→PC synapses throughout CTX, in contrast to hippocampal PV/SST-IN→PC synapses.

Prior research into the effects of systemic morphine administration on PrL observed microcircuit alterations distinct from our acute DAMGO findings (Fig. 3M), with morphine inducing weakened μOR-dependent PV-IN→PC and strengthened δOR-dependent SST-IN→PV-IN inhibition^61^. The surprising δOR-mediated increase in SST-IN output was linked to an upregulation of *Rac1* and *Arhgef6* specifically in SST-INs. While it is unclear if these downstream plasticity mechanisms occur in our acute DAMGO/DPDPE administrations, the high expression of δORs within neocortical PV-INs/SST-INs and inhibitory interconnections between these populations could contribute to the highly variable neocortical opioid modulation we observed.

### Opioids selectively suppress hippocampal synaptic inhibition in nonhuman primates and resected human tissue

To determine if μOR-mediated disinhibitory motifs are present in primate species, we turned to the study of rhesus macaques and resected human tissue. ISH RNAscope analysis of adult macaque tissue revealed that, consistent with mice, the percentage of *Pvalb*+ cells co-expressing *Oprm1* was strongly enriched in CA1 (85.4 ± 4.7%), in contrast to neighboring temporal cortex (16.0 ± 5.3%, Fig. 5A-B). To functionally target PV-INs in macaques, we adopted a viral enhancer strategy. Adult macaques were injected with *pAAV(PHP.eB)-S5E2-ChR2-mCherry* in HPC or M1, an enhancer targeting PV-INs ^62^, which we found to be strongly colocalized with stained PV-INs (Fig. S6A-B). Consistent with our observed PV overlap, S5E2 leIPSCs in acute macaque slices were insensitive to the N-type Ca^2+^ channel blocker ω-conotoxin (95 ± 7%, Fig. S6C), robustly suppressed by the P/Q-type Ca^2+^ channel blocker ω-agatoxin (10 ± 4%, Fig. S6D), and unaffected by the synthetic cannabinoid agonist WIN 55212-2 (97 ± 5%, Fig. S6E), indicating that primate PV-INs utilize similar synaptic mechanisms as in mice. We next assessed the effect of µOR drugs on leIPSCs (Fig. 5C-D). As observed in mice, CA1 leIPSCs were reversibly suppressed by DAMGO (74 ± 3%, Fig. 5E). CA3 leIPSCs exhibited a long-lasting though non-significant DAMGO suppression (66 ± 9%), which was not easily reversed with CTAP or wash (Fig. 5F). DAMGO elicited a highly variable response with no significant suppression in both DG (102 ± 4%, Fig. 5G) and M1 (111 ± 7%, Fig. 5H).

**Fig. 5:**
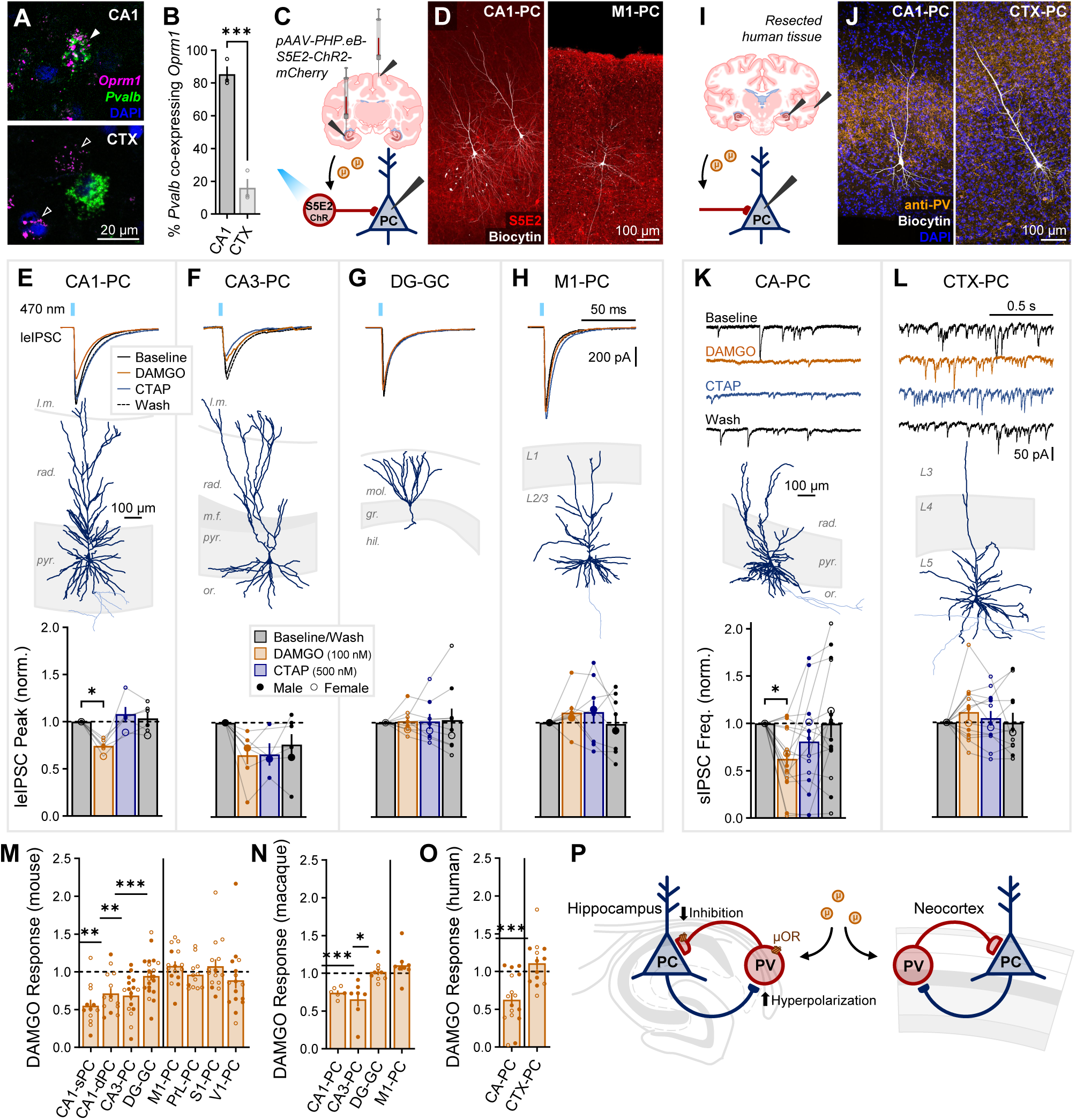
Opioids selectively suppress hippocampal inhibition in nonhuman primates and resected human tissue. (**A**) RNAscope for *Oprm1* and *Pvalb* in adult rhesus macaque. Closed arrow (CA1) indicates *Oprm1*+*Pvalb*+ colocalized cell, while open arrows (CTX) indicate *Oprm1*+*Pvalb*-cells. (**B**) Quantification of the percentage of *Pvalb*+ somata co-expressing *Oprm1* for n_section_ = 3 from 1F, age = 17.6 years. Asterisks represent significance of unpaired t-test. (**C**) Schematic of whole-cell recordings of PCs from S5E2-ChR injected macaques to record the effect of DAMGO/CTAP on leIPSCs. (**D**) Representative *post hoc* staining of recorded CA1 and M1 PCs. (**E**-**H**) Macaque leIPSC experiments with (*top*) averaged example traces, (*middle*) representative *post hoc* reconstructions, and (*bottom*) summary data in (**E**) n_cell_ = 6 CA1-PCs, (**F**) 7 CA3-PCs, (**G**) 9 DG-GCs, and (**H**) 8 M1-PCs, from n_primate_ = 5 (2F), age = 7.4-15.9 (11.2 ± 1.4) years. (**I**) Schematic of whole-cell recordings of PCs from resected human slices to record the effect of DAMGO/CTAP on spontaneous IPSCs (sIPSCs). (**J**) Representative *post hoc* staining of a recorded CA1 and medial temporal cortex PC. (**K**-**L**) Human sIPSC experiments with (*top*) example traces, (*middle*) representative *post hoc* reconstructions, and (*bottom*) summary data in (**K**) n_cell_ = 16 CA-PCs (CA1 & CA3) and (**L**) n_cell_ = 13 CTX-PCs, from n_human_ = 4 (2F), age = 37.3-54.8 (44.8 ± 3.7) years. Asterisks in (*E-L)* represent Tukey’s/Dunn’s *post hoc* comparisons after a significant effect of treatment was found via 1-way repeated measure ANOVA/mixed model/Friedman’s test. (**M**-**O**) Comparison of DAMGO responses across species revealed a similar hippocampal-neocortical divergence in the opioid-mediated suppression of inhibition across (**M**) mice (data from Fig. 3H*-O*, *bottom*), (**N**) macaques (data from *E-H, bottom*), and (**O**) humans (data from *K-L, bottom)*. Asterisks represent significant deviations from the normalized baseline via 1-sample t-test/Wilcoxon. (**P**) Proposed model: hippocampal PV-INs are selectively enriched in µORs, leading to hyperpolarization and suppressed synaptic release.

We next assessed the effect of DAMGO/CTAP on resected human tissue from patients with drug-resistant epilepsy (Fig. 5I). We recorded spontaneous IPSCs (sIPSCs) in CA-PCs (CA1 & CA3) and temporal cortex PCs (Fig. 5J). Although sIPSCs include GABAergic currents from all inhibitory interneurons, perisomatic-targeting FS PV-INs are expected to be overly represented in this measure. Of note, several resections exhibited hippocampal sclerosis within CA1, observable as a marked shrinking of *str. pyr.* with few PCs available for patch clamp electrophysiology. Such sclerosis was likely related to the site of epileptogenesis, and sclerotic resections were not included for this study. Neighboring temporal cortical tissue was surgically removed to reach hippocampal structures and was not pathological. Notwithstanding these limitations, human sIPSCs exhibited a markedly similar regional divergence in DAMGO-mediated suppression, with a significant suppression in CA-PCs (63 ± 8%, Fig. 5K) and no suppression of CTX-PCs (111 ± 8%, Fig. 5L). Thus, although we cannot rule out that pathological activity was present in our human hippocampal resections, the remarkable similarity to healthy murine and nonhuman primate tissue would suggest that opioid-mediated suppression of inhibition was at least unimpaired. Comparing across species, this hippocampal-neocortical divergence was remarkably well-conserved across mice (Fig. 5M), macaques (Fig. 5N), and humans (Fig. 5O). An emerging model for this observed difference is that hippocampal PV-INs are enriched in μORs relative to neocortex, resulting in opioid-mediated hyperpolarization and presynaptic suppression of neurotransmission (Fig. 5P).

### Tac1 cells, as a proxy for immature PV-INs, are suppressed by µOR agonists in HPC

We next explored whether PV-INs exhibit regional opioid divergence in early development. PV-IN function has traditionally been difficult to study in early development as PV itself is not well-expressed until P10 in mice^63^. We undertook an alternate approach from PV^Cre^ mice, with the observation that tachykinin precursor 1 (Tac1, which with post-translational modification produces substance P and neurokinin A) is co-expressed in and can be used as a marker for immature PV-INs^64–67^. To confirm the utility of Tac1 as a marker for immature PV-INs, we examined via snRNAseq of hippocampal interneurons from P10 mice, the expression of principal markers delineating interneurons (Fig. 6A). MGE-INs were distinguished from CGE-INs via expression of *Lhx6*, while *SST* and *Lamp5* (labeling neurogliaform/Ivy cells) labeled clusters of MGE-INs. *Pvalb* at this age was expressed at low levels with high overlap with Tac1. Amongst MGE subgroups at this developmental stage, the future PV cluster was enriched in *Tac1* relative to SST and Lamp5 clusters (Fig. 6B). RiboTag evaluation confirmed that Tac1 associated with the translational machinery was enriched in both HPC and CTX relative to bulk tissue, and in HPC as early as P5 (Fig. 6C).

**Fig. 6:**
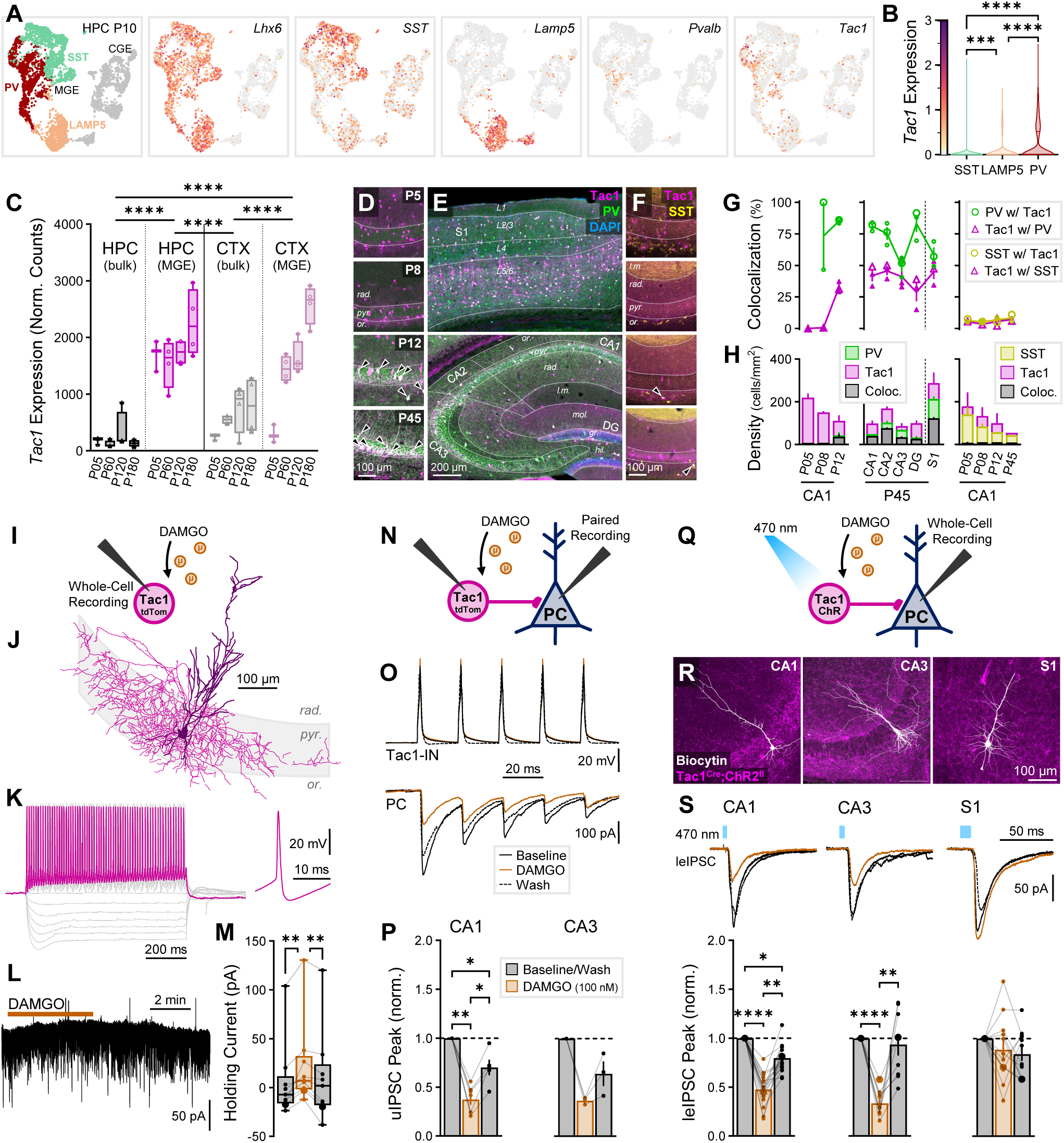
Tac1 cells, as a proxy for immature PV-INs, are suppressed by µOR agonists in HPC. (A) snRNAseq of GABAergic INs in P10 mice across HPC and CTX. *Far left*, Cardinal clusters of *Lhx6*-expressing MGE-INs are colored and delineated as putative SST (*aquamarine*), PV (*red*), and LAMP5 (*peach*) in contrast to caudal ganglionic eminence (CGE)-derived INs (*gray*), with (*left to right*) individual expression of markers *Lhx6*, *SST*, *Lamp5*, *Pvalb*, and *Tac1* indicated. Expression colormap shown in *B*. (B) *Tac1* expression of cardinal MGE-IN cell clusters SST, Lamp5, and PV, n_cell_ = 1476 (SST), 920 (Lamp5), 1451 (PV) from n_mice_ = 2 (1F), age = P10. Asterisks represent results of Dunn’s *post hoc* comparisons after a significant effect of cell type was find via Kruskal-Wallis. (**C**) RiboTag-associated *Tac1* expression in n = 3 P5 and 4 (2F) P60, P120, and P180 Nkx2.1^Cre^: Rpl22(RiboTag)^HA/HA^ mice, comparing bulk HPC/CTX tissue to MGE-INs. Asterisks represent Tukey’s *post hoc* comparisons after significant effects of cell type and region × cell type interaction were observed via 2-way ANOVA. (**D-E**) IHC stain for PV in Tac1^Cre/+^:tdTom^fl/+^ mice, ranging from ages (**D**, *top to bottom*) P5, P8, P12, P45 in CA1, showing the developmental onset of PV expression and colocalization with Tac1, as well as (**E**) in P45 adults throughout S1 cortex (*top*) and HPC (*bottom*). (**F**) IHC stain for SST in Tac1^Cre/+^:tdTom^fl/+^ mice, same ages as *D*. Arrows indicate colocalized cells. Quantification of (**F**) percent colocalization and (**H**) cell density. *Left*: PV-Tac1 CA1 quantification, n_section_ = 2-4 from n_mice_ = 3 each P5, P8, P12. *Middle*: PV-Tac1 quantification for HPC and S1, n_mice_ = 3F P45 (2-4 sections each). *Right*: SST-Tac1 CA1 quantification, n_section_ = 2 from n_mice_ = 2 each P5, P8, P12, P45. (**I**) Schematic of whole-cell recordings of Tac1-INs from P5-11 Tac1^Cre/+^:tdTom^fl/+^ mice to measure intrinsic parameters and the effect of DAMGO. (**J**) Example morphological reconstruction of Tac1-INs (see *Fig. S8*). (**K**) Representative firing of Tac1-IN (see *Fig. S9* and *Table S2*). (**L**) Example trace showing slow currents from DAMGO administration and rapid sEPSCs/sIPSCs. (**M**) Holding current summary data across baseline, DAMGO, and wash conditions for n_cell_ = 9 hippocampal Tac1-INs from n_mice_ = 4 (P8-11). (**N**) Schematic of Tac1→PC paired recordings from P10-11 Tac1^Cre/+^:tdTom^fl/+^ mice to measure the effect of DAMGO on unitary currents (uIPSCs). (**O**) Representative paired recording trace, with repetitive current injection to Tac1-IN to elicit firing (*top*) and record uIPSCs in downstream PC (*bottom*). (P) Paired recording summary data of uIPSC peak amplitude in n_pairs_ = 7 CA1 and 3 CA3 Tac1→PC pairs from n_mice_ = 3 (P10-11). (**Q**) Schematic of whole-cell recordings of PCs recorded in Tac1^Cre/+^:ChR2^fl/+^ mice to record the effect of DAMGO on leIPSCs. (**R**) Representative *post hoc* stains of PCs recorded in CA1, CA3, and S1. (**S**) leIPSC experiments were performed in n_cell_ = 17 CA1-PCs from n_mice_ = 3 (P5-7), n_cell_ = 9 CA3-PCs from n_mice_ = 2 (P6-7), and n_cell_ = 12 S1-PCs from n_mice_ = 3 (P7-10), with (*top*) averaged example traces and (*bottom*) summary data. Asterisks in (*M, P, S*) represent Tukey’s/Dunn’s *post hoc* comparisons after a significant effect of treatment was found via 1-way repeated measure ANOVA/mixed model/Friedman’s test.

We next conducted IHC for PV in P5, P8, P12, and P45 Tac1^Cre/+^:tdTom^fl/+^ mice (Fig. 6D-E). As previously reported, PV was minimally expressed in P5-8 mice but was detectable within CA1 somata and perisomatic axonal terminals by P12. In contrast, Tac1 somata were readily detectable at all ages, with perisomatic axonal terminals detectable and highly colocalized with PV by P12 (Fig. S7A). We quantified somatic colocalization by counting PV+ and Tac1+ cells across hippocampal and neocortical (S1) layers, observing that the percentage of PV-INs co-expressing Tac1 rapidly increased to 85 ± 1% by P12 (Fig. 6G, left), coinciding with the emergence of labeled PV-INs (Fig. 6H, left). CA1 PV co-expression with Tac1 was maintained in adulthood at 81 ± 2% (Fig. 6G, middle). Across subregions and layers, most PV-INs co-expressed Tac1 in HPC (70 ± 4%) and S1 (58 ± 6%). The inverse measure, the percentage of Tac1 cells co-expressing PV, was somewhat lower, rising to ∼50% throughout HPC and S1 in adulthood. Some putative Tac1+ PCs were observed and readily distinguished by morphology and bright fluorescent labeling of dendrites, perhaps representing a limitation of Tac1^Cre/+^:tdTom^fl/+^ mice as they appeared clustered in irregular patches in a minority of sections (27/67 across all ages). Putative Tac1_PCs_ represented a minority fraction, appearing in principal cell layers and comprising 5.9% of all Tac1+ labeled cells. However, their presence warrants caution if employing Tac1^Cre^ mice for functional studies of immature PV-INs without isolating GABAergic transmission or adopting an intersectional approach. Additionally, as there is a reported subpopulation of Tac1-expressing SST-INs^68^, we also stained for SST (Fig. 6F). We observed only a small minority of SST cells co-expressing Tac1 and Tac1 cells co-expressing SST (4-8%, Fig. 6G-H, right, Fig. S7B), consistent with the limited expression of Tac1 in the SST cluster of our snRNAseq data and prior single cell evaluation^69,70^.

We also assessed Tac1-PV colocalization in adult Tac1^Cre/+^:ChR2^fl/+^ mice, observing that across all layers of CA1 the percentage of PV cells co-expressing Tac1 was 48 ± 3%, while the percentage of Tac1 cells co-expressing PV was 74 ± 3% (Fig. S7C-D). We did not observe putative Tac1_PCs_ in this cohort, indicating it may be a reporter-specific observation. These colocalization ratios were effectively inverted from the Tac1^Cre/+^:tdTom^fl/+^ colocalization, likely pointing to the limitations of IHC. ChR2 is a poor somatic label even with immunoboosting, and as PV levels themselves are dynamically regulated^71,72^, several PV-INs likely express sub-threshold levels of PV. In a third IHC cohort, we confirmed that hippocampal Tac1 cells were enriched in μOR receptors by triple labeling Tac1^Cre/+^:μOR-mCherry mice injected perinatally with Cre-dependent *pAAV(AAV5)-pCAG-FLEX-EGFP-WPRE* and staining against PV (Fig. S7E) at P13 and P45. Tac1+PV+ at both ages were >50% colocalized with μORs, while Tac1+PV-were non-significantly less colocalized with μORs (∼40%). However, in total Tac1+ were enriched in μORs relative to Tac1-cells (Fig. S7F).

To more directly assess Tac1 cells as a method to target immature PV-INs, we recorded intrinsic parameters from 90 Tac1+ cells from P5-11 Tac1^Cre/+^:tdTom^fl/+^ mice (Fig. 6I), including from CA1 (44), CA2 (10), CA3 (22), DG (5), and CTX (8). From these cells, 50 hippocampal Tac1+ cells were morphologically reconstructed with visible axon, of which 33/50 (66%) primarily targeted *str. pyr.* and *gr.* consistent with BCs, 11/50 (22%) targeted both *str. or.* and *rad.* consistent with BSCs, 6/50 (12%) primarily targeted superficial dendritic layers, and none exhibited PC morphology (Fig. 6J, Fig. S8). Notably, hippocampal Tac1-IN somata were primarily targeted within *str. pyr*. For comparison of electrophysiological properties, we also recorded from 14 non-labeled putative PCs in HPC, observing striking differences in spiking properties of Tac1-INs versus PCs, consistent with a fast-spiking phenotype expected from PV-INs (Fig. 6K, Fig. S8A-D). We did not observe any regional differences between Tac1-INs (Table S2). To characterize the development of these cells we also recorded from an additional 13 P16-18 Tac1-INs, observing maturation to a faster spiking phenotype similar to 10 PV-INs recorded from mature PV-tdTom mice (Fig. S8E-H). Overall, the developmental trajectory of recorded Tac1-IN physiology and morphology closely matched reported progress of immature to mature PV basket cells^73^.

To assess the opioid sensitivity of Tac1 cells, we applied 100 nM DAMGO to 9 of the recorded hippocampal Tac1 cells in P8-11 mice, observing a reversable hyperpolarizing change in holding current (ΔV = +17 ± 4 pA, Fig. 6L-M). Next, we performed paired recordings of 7 CA1 and 3 CA3 Tac1→PC pairs (Fig. 6N-O) in P10-12 mice, with connected pairs comprising 15% of attempted pairs, observing a strong suppression of transmission in both regions (CA1: 37 ± 5%, CA3: 37 ± 2%, Fig. 6P) and consistent with adult PV→PC pairs^21^. Utilizing Tac1^Cre/+^:ChR2^fl/+^ mice (Fig. 6Q-R), we further explored the opioid modulation of inhibitory output across regions. In P5-7 mice we observed that GABAergic-isolated leIPSCs were reversibly suppressed by DAMGO in both CA1 (47 ± 4%) and CA3 (34 ± 4%, Fig. 6S). Within S1-L5, we observed that functional Tac1→PC light-evoked synaptic responses were often small and unreliable before the age of ∼P8, thus we included in our study slightly older mice with a full age range from P7-10. Consistent with adult PV-INs, we did not observe a significant suppression of Tac1 mediated inhibition (89 ± 10%, Fig. 6S). Thus, at the earliest timepoints just as immature PV-IN→PC synaptic connections are being established, there is opioid receptor dependent modulation within the hippocampus not found in neocortex.

### Opioids and Tac1 cells regulate spontaneous activity of the developing hippocampus

During early postnatal development between P5-10, the principal network signature is that of giant depolarizing potentials (GDPs). These spontaneous synchronous events occur during this critical developmental window while GABA is depolarizing^74^ and are believed to play an important role in hippocampal synaptogenesis^75^. To test the importance of opioids in regulating these events, in P5-8 mice we recorded spontaneous GDP associated currents (GDP-Is) intracellularly from CA3-PCs voltage-clamped to 0 mV to isolate GABAergic contributions, in which GDP-Is are detectable as outward events due to a barrage of GABA_A_ currents (Fig. 7A-B). GDP-I rate was highly variable across brain slices, so to ensure enough events for averaging we only considered slices with ≥ 4 baseline GDPs (69/105 slices across all conditions). We next applied 100 nM DAMGO and observed a robust and reversible decrease in GDP-I event frequency (32 ± 6% of baseline), with no change in GDP-I amplitude (Fig. 7C). DAMGO also suppressed GDPs within S1 cortex (21 ± 9%), which we observed occurring at a lower rate than in CA3 (Fig. S10A-B). δOR agonist DPDPE could also suppress CA3 GDPs (55 ± 13%, Fig. S10C-D), while µOR antagonist CTAP had no effect (104 ± 16%, Fig. S10E-F). Thus, opioid agonists potently suppress spontaneous network activity of the developing brain, analogous to the effect of DAMGO on spontaneous sharp wave ripples (SWRs) in adulthood^41^. To determine if Tac1 cells could subserve this suppressing role, we optogenetically silenced Tac1 cells with Tac1^Cre/+^:ArchT^fl/+^ mice (Fig. 7D-E), observing that GDP-Is were significantly suppressed (76 ± 8% of baseline, Fig. 7F), with no effect on sIPSCs (Fig. S10G). Moreover, Tac1 cell silencing occluded the effect of DAMGO, as DAMGO produced no further reduction in GDP-I frequency (97 ± 16% compared to light-on period, Fig. 7G-I). Prior work from our lab identified MGE-INs as key regulators of GDP activity; optogenetically silencing MGE-INs reduces GDP-I frequency to 34 ± 6% of baseline^76^. Several studies have attributed GDP regulation and generation to dendritic-targeting SST-INs^77,78^. However, optogenetically silencing SST-INs results in only modest reductions of GDP-I frequency (71-80% of baseline)^79^. Thus, to account for the entire effects of DAMGO-mediated suppression and optogenetic MGE-IN silencing, other IN subpopulations likely contribute. This study presents for the first-time evidence that immature PV-INs play a key role in early hippocampal population activity, consistent with the extensive and functional perisomatic innervation already provided by these cells at this stage of development (Fig. 6I-P, Fig. S8) and the essential role adult PV-INs play in hippocampal rhythmogenesis. The limited information regarding immature PV-IN participation in developmental hippocampal population activity likely reflects a prior lack of viable animal model to target these cells. Importantly, the suppression of immature PV-INs by opioids at these early developmental time-points has severe implications for the harmful aspects of opioid use on synaptogenesis and circuit development in the developing brain.

**Fig. 7:**
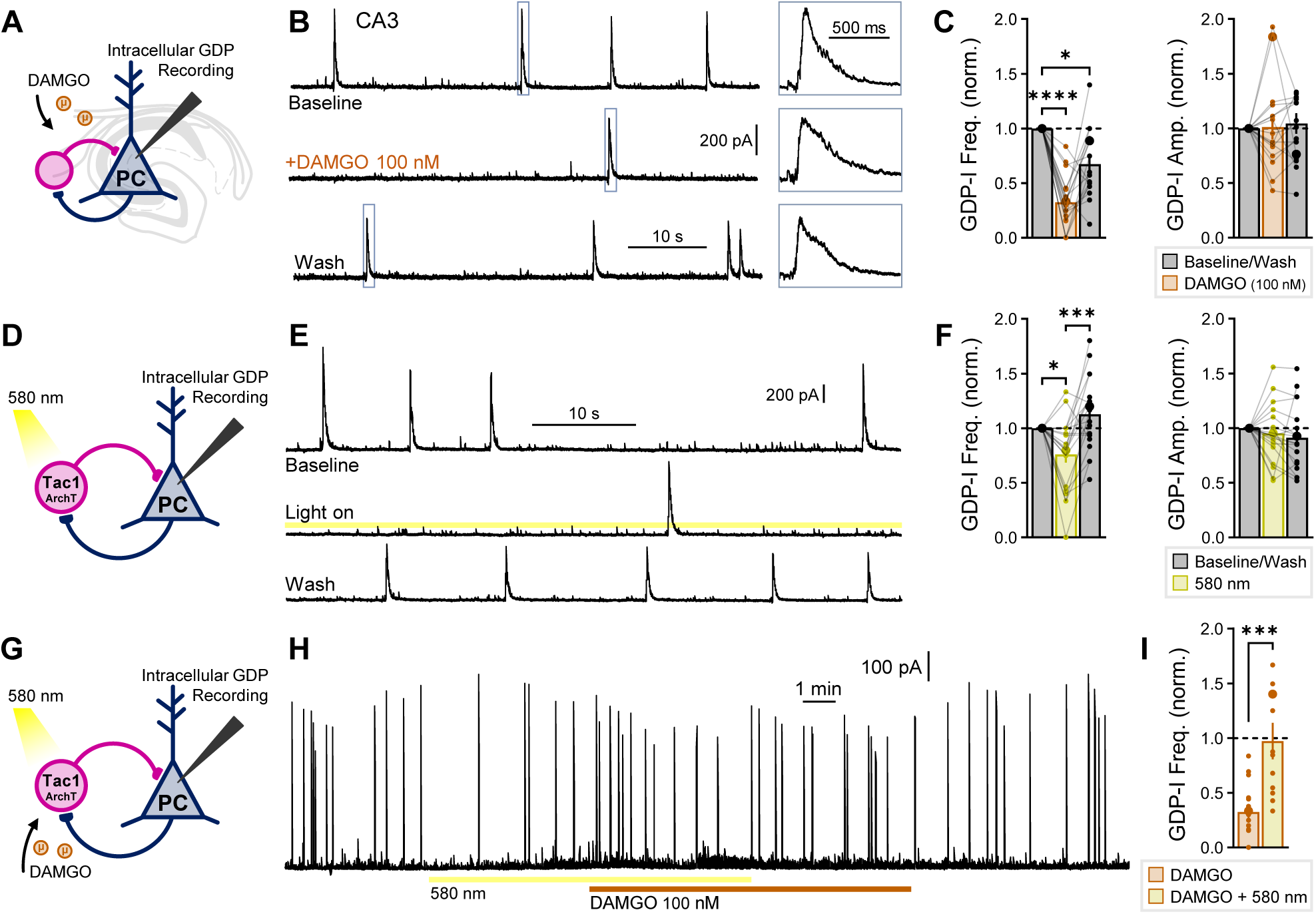
Opioids and Tac1 cells regulate spontaneous activity of the developing hippocampus. (**A**) Schematic of recordings of GDP associated currents (GDP-Is), recorded intracellularly in CA3-PCs voltage-clamped to 0 mV in WT mice. (**B**) Example traces of the effect of 100 nM DAMGO applied for 5 minutes, with (*right inset*) example GDP-I events. (**C**) DAMGO summary data for GDP-I event frequency and amplitude from n_cell_ = 18 from n_mice_ = 6 (P5-8). (**D**) Schematic of CA3-PC GDP-I recordings from Tac1^Cre/+^:ArchT^fl/+^ mice to silence Tac1 cells. (**E**) Example traces of the effect of ArchT activation with 580 nm light for 1-2 min. (**F**) ArchT summary data for GDP-I event frequency and amplitude from n_cell_ = 20 from n_mice_ = 6 (P5-8). Asterisks in (*C*, *F*) represent Dunn’s *post hoc* comparisons after a significant effect of treatment was found via Friedman’s test. (**G**) Schematic of ArchT-DAMGO occlusion in CA3-PCs. (**H**) Example trace of the effect of Tac1-ArchT silencing for 10 min., with 100 nM DAMGO applied 5 min. after start of Tac1-ArchT silencing. (**I**) ArchT-DAMGO occlusion summary data from n_cell_ = 13 from n_mice_ = 4 (P6-8). DAMGO administration following Tac1-AchT silencing did not further suppress GDP-I event frequency (*yellow*). DAMGO alone (*orange*) plotted from *C* for comparison. Asterisks represent results of unpaired t-test.

### Morphine pre-treatment occludes acute DAMGO suppression

Exposure to exogenous opioids can result in a shift in μOR/δOR function to become more constitutively active in chronic^80^ and withdrawal conditions^59,81^, which is believed to contribute to tolerance and dependence. Drug-induced alterations to the endogenous opioid system have not been well-studied in the hippocampus. One study found chronic morphine treatment results in altered RNA levels of *Penk* and *Gnas* (encoding the G_αs_ subunit of GPCRs)^82^, although it is unclear if these changes are specific to distinct neuronal populations. Therefore, to determine if the opioid-mediated suppression of PV-INs is altered after morphine exposure, we injected morphine or saline in PV^Cre/+^:ChR2^fl/+^ mice prior to acute slice preparation and whole cell patch clamp recordings of leIPSCs. We used three treatment regimens (Fig. 8A), each with an independent saline control group, including a bolus injection (15 mg/kg), a chronic treatment with increasing doses over 6 days (15, 20, 25, 30, 40, 50 mg/kg), and a withdrawal group receiving the chronic treatment but with acute slice preparation 72 hours instead of 1 hour after the final injection as with other groups. Optogenetic leIPSC recordings were performed in CA1-PCs as previously described. Saline controls all exhibited significant DAMGO-mediated suppression of leIPSCs (Fig. 8B) similar to un-injected mice (cf. Fig. 3H-I). In morphine-injected mice, the acute DAMGO-mediated suppression was partially occluded, particularly in the withdrawal group for which DAMGO no longer significantly deviated from baseline (Fig. 8C). Comparing the normalized DAMGO response across all groups (Fig. 8D), the bolus and withdrawal groups exhibited a significant deviation from their saline control, which was somewhat moderated in the chronic group, potentially through long-term adaptations distinct from the withdrawal conditions where constitutive receptor activity has been described^59,81^. Although the physiological role of the opioid-mediated suppression of hippocampal PV-INs is not yet fully understood, these data indicate that this disinhibitory motif becomes dysregulated after morphine use. Exogenous opioids, in addition to well-described roles in analgesic and stress response, can therefore interfere with neuromodulation of hippocampal PV-INs, leading to disruption of rhythmic activity such as GDPs (Fig. 7), SWRs^41^, and gamma oscillations^42^, activities essential for learning and memory in the developing and adult brain.

**Fig. 8:**
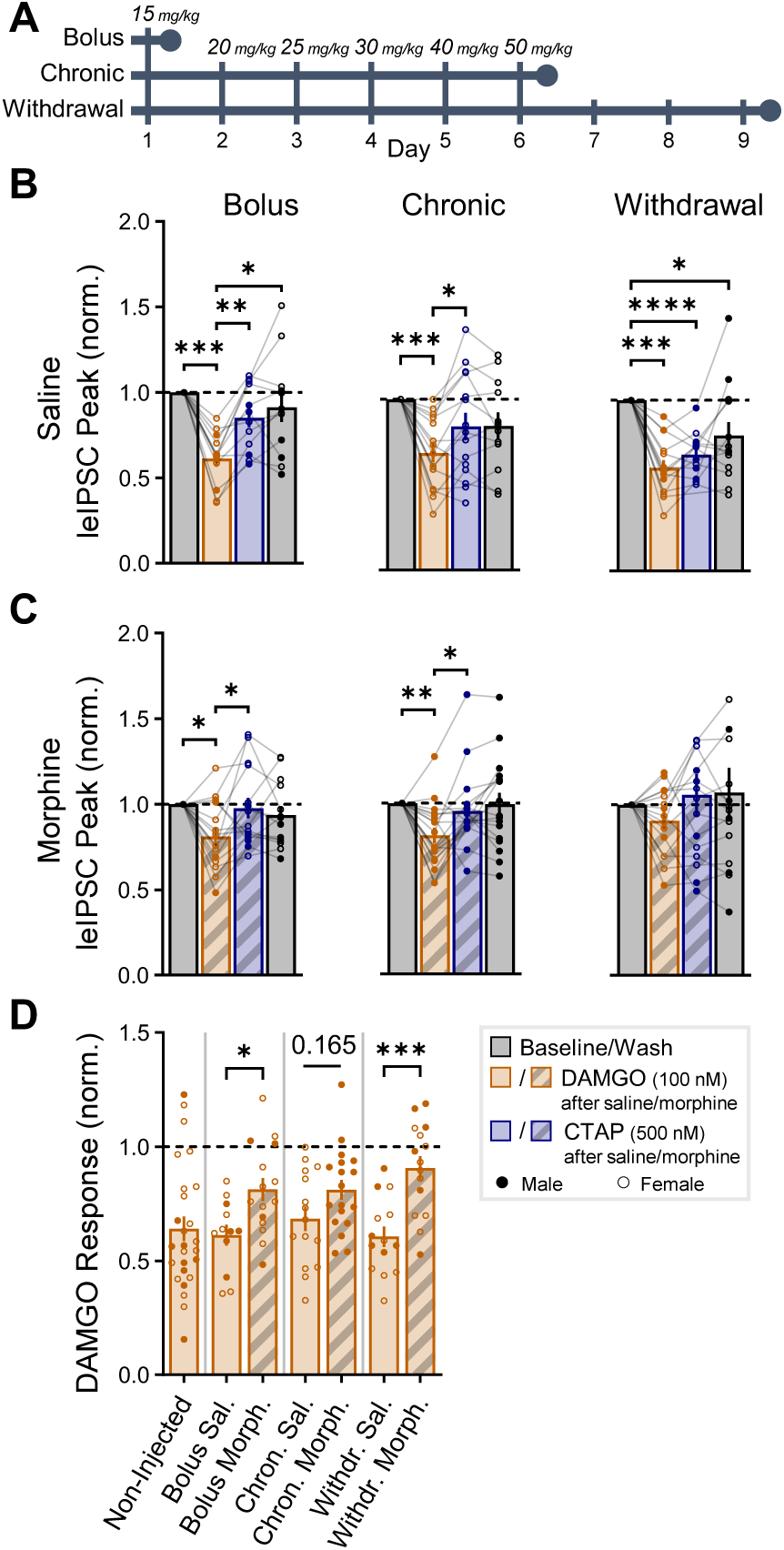
Morphine pre-treatment occludes acute DAMGO suppression. (**A**) Morphine pretreatment experimental design with 3 regimens: bolus (15 mg/kg), chronic (15, 20, 25, 30, 40, 50 mg/kg) and withdrawal (chronic regimen + 72 hours). All injections were performed in adult PV^Cre/+^:ChR2^fl/+^ mice with either daily morphine or saline injections, for a total of 6 groups. (**B**) Saline control groups all exhibited significant DAMGO suppression of leIPSCs recorded in CA1-PCs (cf. Fig. 3H*-I*). *Bolus*: n_cell_ = 12 from n_mice_ = 3 (2F), age = P64-126 (P103 ± 20). *Chronic*: n_cell_ = 15 from n_mice_ = 3F, age = P67-76 (P73 ± 3). *Withdrawal*: n_cell_ = 14 from n_mice_ = 3 (2F), age = P78-83 (P80 ± 2). (**C**) Morphine injected groups exhibited partial occlusion of acute DAMGO-mediated suppression of leIPSCs, particularly in the withdrawal group. *Bolus*: n_cell_ = 15 from n_mice_ = 3 (2F), age = P65-127 (P104 ± 20). *Chronic*: n_cell_ = 18 from n_mice_ = 3M, age = P68-76 (P72 ± 2). *Withdrawal*: n_cell_ = 15 from n_mice_ = 3 (2F), age = P77-81 (P79 ± 1). Asterisks in (*B-C*) represent Tukey’s *post hoc* comparisons after a significant effect of treatment was found via 1-way repeated measure ANOVA/mixed model. (**D**) Combined DAMGO responses (non-injected data from Fig. 3H*-I* for comparison). Asterisks represent Šídák’s *post hoc* comparisons after a significant effect of treatment was found via 2-way ANOVA, with comparisons restricted to those of *a priori* interest (morphine vs. saline for each regimen).

## DISCUSSION

PV-INs are essential gatekeepers of cortical activity, supporting excitatory:inhibitory (E:I) balance, feedforward inhibition, gamma oscillations, SWRs, and as observed in this study, GDPs. The ability of opioids to suppress PV-IN activity has critical implications for these roles. This study demonstrates that outside the hippocampus proper, PV-INs largely do not express µOR receptors and are unaffected by µOR agonists. Consistent with these findings, early micro-iontophoresis studies observed that administering enkephalin throughout CTX suppresses network spiking activity, in contrast to HPC where enkephalin increases network activity via disinhibition^14,18^. IHC studies have revealed that µOR-expressing neocortical cells are overwhelmingly GABAergic (97%), but primarily vasoactive intestinal peptide-expressing interneurons (VIP-INs, 92%) rather than PV-INs (8%)^83^. Intriguingly, neocortical µOR-expressing cells largely co-express *Penk*, the precursor to enkephalin, suggesting an auto-suppressing role of these cells. Within HPC, VIP-INs are also the primary source of enkephalin^84^, the release of which supports plasticity and social memory^85^. Thus, VIP-INs are the local source of endogenous opioids in both HPC and CTX, but the downstream receptor-expressing cells have shifted to include PV/SST-INs in HPC.

Contrasting with this dichotomy, some studies have found evidence for opioid-mediated suppression of PV-INs in select regions of neocortex. In the medial entorhinal cortex (mEC), µ-opioids suppress GABA release from fast-spiking (FS) PV-INs onto stuttering PV-INs, with a consequent increase in gamma oscillations^86^. Voltage-dependent Na^+^ currents in prefrontal GABAergic INs are suppressed by DAMGO, though this study did not delineate GABAergic subtype^87^. Within insular cortex, paired recordings from FS, non-FS, and PCs indicate that DAMGO suppresses FS→FS, non-FS→FS but not FS→PC, non-FS→PC connections, while DPDPE suppresses FS→FS, FS→PC but not non-FS→FS, non-FS→PC connections^88^, pointing to a complex interplay of interneuronal opioid modulation. Notably, these cortical findings largely agree with our own, with no effect of DAMGO on PV-IN→PC, DAMGO/DPDPE on SST-IN→PC, and a subthreshold effect of DPDPE on PV-IN→PC connections. In orbitofrontal cortex (OFC), electrical- and light-evoked PV-IN inputs to PCs display sensitivity to DAMGO in medial (m) but not lateral (l)OFC^89^. Within neighboring prelimbic cortex (PrL), systemic morphine suppresses light-evoked PV-IN release via μORs but increases SST-IN release onto PV-INs via δORs^61^. Although we observed no significant DAMGO-mediated suppression of PV-IN release across cortex (including PrL), the highly variable responses observed and complex interplay of interneuronal circuitry across CTX evidenced in these studies may prove distinct from microcircuitry motifs in HPC. Another possibility is that phylogenetically older cortical regions such as allocortical HPC, mEC, PrL, and mOFC^90^ exhibit region-specific transcriptomic, translational, or post-translational programs permitting the expression of functional μORs within PV-INs.

While we observed no evidence that earlier-born PV-INs expressed more μORs (Fig. S2), we did replicate observations of a shifted developmental period of neurogenesis within HPC (E11.5^43^) versus CTX (E13.5^47,48^). In our Tac1 experiments we observed that functional neocortical Tac1→PC synaptic connections were delayed relative to HPC, becoming prominent around P8, while within HPC they were robustly observed at the earliest ages studied (P5). Within HPC, early-born E11.5 PV-INs are reported to express higher PV levels, receive higher synaptic E:I ratios, and preferentially innervate dPCs over sPCs in contrast to late-born E13.5 PV-INs^72^. Within the SST-IN population, a subpopulation of early-born hub cells critically regulate GDPs^77,78^. Thus, the birth date of interneurons strongly determines circuit connectivity and function and, as has been suggested^91^, may establish an additional criterion to classify neuronal populations.

The energy demands of maintaining functional opioid receptors in hippocampal PV-INs suggest they support important physiological roles. Both exogenous and endogenous opioid release promote long-term potentiation (LTP) within the lateral perforant path to DG^92,93^. Within CA1, LTP is altered in rats chronically treated with morphine^94,95^. The specific PC sub-compartments innervated by opioid-sensitive inputs could also have profound effects on dendritic integration. Although we observed dendritic-targeting CA1 SST-INs were suppressed by opioids (Fig. 4K), as well as prior work demonstrating NPY-INs are suppressed by opioids^27^, perisomatic-targeting PV-INs may be the greatest contributors to opioid-mediated disinhibition of PCs. In addition to expressing the highest levels of µORs^24,25^, perisomatic inhibition targeting *str. rad*., *pyr*. and *or*. is more susceptible to opioid suppression relative to distal *l.m.*^96^, suggesting that opioid-mediated disinhibition may bias a PC to preferentially integrate more proximal Schaffer collaterals relative to distal *l.m.* inputs. Opioids may also play a key role in place field formation. During theta oscillations, a transient reduction in inhibition is critical for place field formation^97^. More specifically, morphine causes place field remapping to environments associated with drug administration^98^. Future studies are needed to explore how place field formation, learning, and memory consolidation are affected by selective PV-IN modulation and opioid treatments.

The present study establishes that opioids are uniquely positioned to suppress hippocampal proper and not primary neocortical PV-INs. In contrast to reports of PV-IN evolutionary divergence^39,40^, our data demonstrate this disinhibitory circuit motif is maintained across mice, macaques, and humans. This motif is established in early development, and both opioids and immature PV-INs critically regulate hippocampal population activity, suggesting one potential mechanism for neurodevelopmental deficits due to chronic *in utero* opioid exposure^4^. These findings highlight that opioids acting through immature PV-INs have an underappreciated role in regulating developmental hippocampal activity.

## Supporting information

Table S1: Statistical details

Table S2: Intrinsic Parameters

## ACKNOWLEDGEMENTS

This work was supported by a NICHD Intramural Research Program (IRP) grant to CJM, NINDS IRP grant to KZ, NIMH IRP grant to BBA, and NIH Center on Compulsive Behaviors (CCB) fellowship to APC. This work was carried out in collaboration with the NIH Comparative Brain Physiology Consortium (CBPC) at the NIH IRP. RNA sequencing and analysis support was provided by the Molecular Genomics Core, Bioinformatics and Scientific Programming Core, NICHD. Imaging support was provided by Vincent Schram at the NICHD Microscopy and Imaging Core.

## AUTHOR CONTRIBUTIONS

APC was responsible for project conceptualization, experimental design, electrophysiology experiments, imaging, analysis, and principal writer of this manuscript. KAP and CJM conceptualized, designed, and supervised the project. JHK, GV, LH, RC, and KAP contributed to electrophysiology experiments. AV, NM, SK, XY, SH, AR, and IS contributed to IHC and ISH. VM conducted RiboTag experiments and analysis. EL assisted with morphine injections. EF, MD, BLG, and GF conducted snRNAseq data collection, analysis, and interpretation. DA supported mouse breeding, colony maintenance, and genotyping. MAGE, ACC, BEH, AP, AM, and BBA conducted NHP handling, injections, and surgery. KZ conducted surgical procedures with human patients. GF and JD provided support and design of viral constructs. All authors participated in drafting the manuscript. MAGE, BBA, KZ, and CJM provided critical resources for the study.

## DECLARATION OF INTERESTS

The authors declare no competing interests.

## METHODS

### KEY RESOURCES TABLE

| REAGENT or RESOURCE | SOURCE | IDENTIFIER |
| --- | --- | --- |
| Antibodies |  |  |
| Rabbit anti-Parvalbumin | Abcam | Cat# ab11427, RRID:AB_298032 |
| Mouse anti-Parvalbumin | Sigma | Cat# P3088, RRID:AB_477329 |
| Rabbit anti-Somatostatin 28 | Abcam | Cat# ab111912, RRID:AB_10903864 |
| Rabbit anti-μOR | Abcam | Cat# ab134054, RRID:AB_3122135 |
| Rabbit anti-Satb1 | Abcam | Cat# ab109122, RRID:AB_10862207 |
| Rabbit anti-plkBα | Cell Signaling Technology | Cat# 2859, RRID:AB_561111 |
| Chicken anti-GFP | Aves Labs | Cat# GFP-1020, RRID:AB_10000240 |
| Rabbit anti-GFP | Abcam | Cat# ab290, RRID:AB_303395 |
| Guinea Pig anti-RFP | Synaptic Systems | Cat# 390005, RRID:AB_2737051 |
| Rabbit anti-RFP | Antibodies-Online | Cat# ABIN129578, RRID:AB_10781500 |
| AF488 Goat anti-Rabbit IgG (H+L) | Thermo Fisher Scientific | Cat# A-11008, RRID:AB_143165 |
| AF488 Goat anti-Rabbit IgG (H+L) | Thermo Fisher Scientific | Cat# A-11034, RRID:AB_2576217 |
| AF488 Donkey anti-Rabbit IgG (H+L) | Thermo Fisher Scientific | Cat# A-32790, RRID:AB_2762833 |
| CF488A Donkey anti-Mouse IgG (H+L) | Biotium | Cat# 20014, RRID:AB_10561327 |
| CF488A Donkey anti-Rabbit IgG (H+L) | Biotium | Cat# 20015, RRID:AB_10559669 |
| CF488A Donkey anti-Chicken IgY (IgG) (H+L) | Biotium | Cat# 20166, RRID:AB_10854387 |
| AF555 Goat anti-Rabbit IgG (H+L) | Thermo Fisher Scientific | Cat# A-21429, RRID:AB_2535850 |
| AF555 Goat anti-Guinea Pig IgG (H+L) | Thermo Fisher Scientific | Cat# A-21435, RRID:AB_2535856 |
| CF555 Donkey anti-Rabbit IgG (H+L) | Biotium | Cat# 20038, RRID:AB_10558011 |
| CF555 Donkey anti-Guinea Pig IgG (H+L) | Biotium | Cat# 20276-1, RRID:AB_10853597 |
| AF594 Donkey anti-Guinea Pig IgG (H+L) | Thermo Fisher Scientific | Cat# A-11076, RRID:AB_141930 |
| CF647 Donkey anti-Mouse IgG (H+L) | Biotium | Cat# 20046-1, RRID:AB_10853778 |
| CF647 Donkey anti-Rabbit (H+L) | Biotium | Cat# 20047, RRID:AB_10559808 |
| Bacterial and virus strains |  |  |
| pAAV(PHP.eB)-BiPVe4-ChR2-mCherry | Addgene<sup>60</sup> | RRID:Addgene_213937 |
| pAAV(PHP.eB)-BiPVe4-tdTom-tdTom<sub>nls</sub> | Addgene<sup>60</sup> | RRID:Addgene_213936 |
| pAAV(AAV9)-Ef1a-DIO-C1V1(t/t)-TS-mCherry | Addgene | RRID:Addgene_100061-AAV9 |
| pAAV(PHP.eB)-S5E2-ChR2-mCherry | Addgene<sup>62</sup> | RRID:Addgene_135634-PHPeB |
| pAAV(AAV5)-pCAG-FLEX-EGFP-WPRE | Addgene<sup>99</sup> | RRID:Addgene_51502-AAV5 |
| Chemicals, peptides, and recombinant proteins |  |  |
| DAMGO | Tocris | Cat# 1171 |
| CTAP | Tocris | Cat# 1560 |
| DPDPE | Tocris | Cat# 1431 |
| Naltrindole hydrochloride | Millipore Sigma | Cat# N115 |
| Morphine Sulfate Injection, USP (4 mg/mL) | Hikma Pharmaceuticals | NDC: 0641-6125 |
| Naloxone Hydrochloride Injection (0.4 mg/mL) | Mylan Institutional | NDC: 67457-292-02 |
| DNQX disodium salt | Tocris | Cat# 2312 |
| DL-APV | Tocris | Cat# 0105 |
| CGP 55845 hydrochloride | Tocris | Cat# 1248 |
| ω-Agatoxin IVA | Tocris | Cat# 2799 |
| ω-Conotoxin GVIA | Tocris | Cat# 1085 |
| WIN 55212-2 mesylate | Tocris | Cat# 1038 |
| Biocytin | Millipore Sigma | Cat# B4261 |
| AF488 streptavidin | Thermo Fisher Scientific | Cat# S11223 |
| AF555 streptavidin | Thermo Fisher Scientific | Cat# S21381 |
| DAPI | Millipore Sigma | Cat# D9542 |
| RNAscope mouse probe Mm-Pvalb | Advanced Cell Diagnostic | Cat# 421931 |
| RNAscope mouse probe Mm-Oprm1-C2 | Advanced Cell Diagnostic | Cat# 315841-C2 |
| RNAscope mouse probe Mm-Oprd1 | Advanced Cell Diagnostic | Cat# 427371 |
| RNAscope macaque probe Mmu-PVALB | Advanced Cell Diagnostic | Cat# 461691 |
| RNAscope macaque probe Mmu-OPRM1-C2 | Advanced Cell Diagnostic | Cat# 518941-C2 |
| EdU | Thermo Fisher Scientific | Cat# A10044 |
| Opal 520 | AKOVA | Cat# OP-001001 |
| Opal 570 | AKOVA | Cat# OP-001003 |
| ProLong Diamond Antifade Mountant with DAPI | Thermo Fisher Scientific | Cat# P36962 |
| ProLong Gold Antifade Mountant | Thermo Fisher Scientific | Cat# P10144 |
| GTPyS-Li<sub>4</sub> | Sigma | Cat# G8634 |
| **Critical commercial assays** |  |  |
| Chromium Single Cell 3' GEM, Library & Gel Bead Kit v3 | 10x Genomics | Cat# PN-1000075 |
| RNAscope Multiplex Fluorescent v2 Assay | Advanced Cell Diagnostic | Cat# 323100 |
| RNeasy Plus Micro Kit | Qiagen | Cat# 74034 |
| RNA 6000 Pico kit | Agilent | Cat# 5067-1513 |
| SMARTer Stranded Total RNA-Seq. Kit v2 | Takara Bio | Cat# 634411 |
| Click-iT EdU imaging Kit | Thermo Fisher Scientific | Cat# C10340 |
| **Experimental models: Organisms/strains** |  |  |
| Mouse: PV<sup>Cre/Cre</sup> | Jax<sup>100</sup> | RRID:IMSR_JAX:017320 |
| Mouse: SST<sup>Cre/Cre</sup> | Jax<sup>101</sup> | RRID:IMSR_JAX:013044 |
| Mouse: Tac1<sup>Cre/Cre</sup> | Jax<sup>102</sup> | RRID:IMSR_JAX:021877 |
| Mouse: Nkx2.1<sup>Cre/Cre</sup> | Jax<sup>103</sup> | RRID:IMSR_JAX:008661 |
| Mouse: Dlx5/6<sup>Cre/Cre</sup> | Jax<sup>104</sup> | RRID:IMSR_JAX:008199 |
| Mouse: tdTomato(Ai9)<sup>fl/fl</sup> | Jax<sup>105</sup> | RRID:IMSR_JAX:007909 |
| Mouse: ChR2(Ai32)-eYFP<sup>fl/fl</sup> | Jax<sup>106</sup> | RRID:IMSR_JAX:024109 |
| Mouse: ArchT(Ai40D)-eGFP<sup>fl/fl</sup> | Jax<sup>107</sup> | RRID:IMSR_JAX:021188 |
| Mouse: Rpl22(RiboTag)<sup>HA/HA</sup> | Jax<sup>44</sup> | RRID:IMSR_JAX:011029 |
| Mouse: CAG-Sun1/sfGFP<sup>fl/fl</sup> | Jax<sup>108</sup> | RRID:IMSR_JAX:030952 |
| Mouse: Oprm1<sup>fl/fl</sup> | Jax | RRID:IMSR_JAX:030074 |
| Mouse: μOR-mCherry | Jax<sup>31</sup> | RRID:IMSR_JAX:029013 |
| Mouse: PV-tdTomato | Jax | RRID:IMSR_JAX:027395 |
| Mouse: C57BL/6J | Jax | RRID:IMSR_JAX:000664 |
| **Software and algorithms** |  |  |
| pClamp 10.7 | Molecular Devices | RRID:SCR_011323 |
| FIJI (ImageJ) | NIH | RRID:SCR_002285 |
| Excel | Microsoft | RRID:SCR_016137 |
| Prism | Graphpad | RRID:SCR_002798 |
| ZEN Microscopy Software | Zeiss | RRID:SCR_013672 |
| SPOT Basic 5.2 | SPOT Imaging | RRID:SCR_014313 |
| CellRanger v7.0.0 | 10x Genomics |  |
| DESeq2 | Bioconductor<sup>109</sup> | RRID:SCR_015687 |
| FIJI Macro: colocIHC | this manuscript | https://doi.org/10.5281/zenodo.10063200 |

### RESOURCE AVAILABILITY

#### Lead Contact

Further information and requests for resources and reagents should be directed to and will be fulfilled by the lead contact, Kenneth Pelkey.

#### Materials Availability

This study did not generate novel, unique reagents.

#### Data and Code Availability

Data generated during this study are available upon request. The custom ImageJ macro developed to automate IHC colocalization is open-source and available via public repositories: a current version subject to change (https://github.com/acaccavano/colocalizationIHC) and an archival copy used for this manuscript (https://doi.org/10.5281/zenodo.10063200).

### EXPERIMENTAL MODEL AND STUDY PARTICIPANT DETAILS

#### Mice

Targeted labeling of neuronal subpopulations was achieved through Cre-recombinase driven expression of floxed reporters/genes, with experimental offspring maintained as heterozygous crosses of the homozygous Cre driver and homozygous floxed reporter mouse lines listed in the Key Resources Table. Ribotag experiments with Nkx2.1^Cre/+^:Rpl22(RiboTag)^HA/+^ offspring were cross-bred for several generations to obtain homozygosity in Rpl22^HA/HA^, to account for the relatively low expression of the endogenous *Rpl22* gene^45,46^. To selectively knock-out *Oprm1* in PV-INs, PV^Cre/+^:Oprm1^fl/+^ offspring were crossed with Oprm1^fl/fl^ mice to achieve homozygosity in floxed *Oprm1*. Viral injections and experiments were also performed in C57BL6/J wild-type (WT) mice.

Male and female mice were used in approximately equal number, with those above weaning age sexed by inspection. The number of experimental mice used for this study were: 71 (36F) PV^Cre/+^:ChR2^fl/+^ P27-144 (P74 ± 3), 29 C57BL6/J P6-155 (P33 ± 6), 22 Tac1^Cre/+^:tdTomato^fl/+^ P5-45 (P14 ± 3), 15 Nkx2.1^Cre/+^:Rpl22(RiboTag)^HA/HA^ P5-180 (P97 ± 17), 13 (8F) PV-tdTomato^+/-^ P37-392 (P136 ± 39), 11 Tac1^Cre/+^:ChR2^fl/+^ P5-37 (P15 ± 4), 9 Tac1^Cre/+^:ArchT^fl/+^ P5-8 (P6.8 ± 0.3), 9 (5F) SST^Cre/+^:ChR2^fl/+^ P57-248 (P146 ± 25), 8 (4F) Tac1^Cre/+^:μOR-mCherry^+/-^ P13-45 (P29 ± 6), 4 (2F) Dlx5/6^Cre^:CAG-Sun1/sfGFP P10-28 (P19 ± 5), 3M PV^Cre/+^:Oprm1^fl/fl^ P58-77 (P65 ± 6), 1F PV^+/+^:Oprm1^fl/fl^ P77, and 2F μOR-mCherry^+/-^ P70. Mice were housed and bred in a conventional vivarium with standard laboratory chow and water in standard animal cages under a 12 hr circadian cycle. All rodent experiments were conducted under an Animal Study Protocol approved by the ACUC at the National Institute of Child Health and Human Development.

#### Nonhuman Primates (NHPs)

Primate tissue was obtained from 7 (4F) adult rhesus macaques, aged 7-17 (12.2 ± 1.3) years, that had reached the end of their paradigms for other experiments, as part of the NIH comparative brain physiology consortium (CBPC). Of these macaques, 6 were virally injected (5 for acute slice preparation and 1 fixed-perfused for IHC) and 1 was un-injected with tissue flash-frozen for ISH RNAscope. All experiments were performed in accordance with the ILAR Guide for the Care and Use of Laboratory Animals and were conducted under an Animal Study Protocol approved by the ACUC at the National Institute of Mental Health. All procedures adhered to the applicable Federal and local laws, regulations, and standards, including the Animal Welfare Act and Regulations and Public Health Service policy (PHS2002).

#### Human Tissue

Human tissue was obtained from surgical specimens collected from 4 (2F) anonymized/deidentified patients with pharmaco-resistant epilepsy, aged 37-54 (44.8 ± 3.7) years. The participants underwent an initial surgical procedure during which recording electrodes were implanted subdurally on the cortical surface and within the brain parenchyma to monitor epileptiform activity. The location of the intracranial electrodes was selected by the clinical team to localize the epileptogenic zone and recordings during a monitoring period were used to identify the specific hippocampal region exhibiting ictal or inter-ictal activity. During a second surgery the brain areas of seizure onset were surgically resected. The Institutional Review Board (IRB) at the National Institute of Neurological Disease and Stroke approved the research protocol and informed consent for the experimental use of surgically resected tissue was obtained from each participant and their guardian.

## METHOD DETAILS

### Viral Injections (mouse)

For retro-orbital injections of *pAAV(PHP.eB)-BiPVe4-ChR2-mCherry* (Addgene 213937) and *pAAV(PHP.eB)-BiPVe4-tdTom-tdTom_nls_* (Addgene 213936) viruses, mice were anesthetized with 5% isoflurane then transferred to a nose-cone delivering 2% isoflurane for the duration of the injection. Mice were injected with 80 μL of virus (1:9 dilution with sterile 1x PBS) with a 0.5 mL insulin syringe (BD Syringes) in the retro-orbital sinus.

For intra-cranial injections, mice were anesthetized with 5% isoflurane and mounted on a stereotax (Neurostar). Mouse were delivered 2% percent isoflurane throughout surgery. Subcutaneous injection of buprenorphine (0.3 mg/ml) and topical application of lidocaine/prilocaine ointment (2.5%/2.5%) were provided for post-operative analgesia. Injections were delivered with a glass micropipette (Neurostar) at a rate of 100 nL/min. Following injection, the pipette was left in place at site for 5 minutes before removal. Each craniotomy was plugged with Kwik-Sil silicone elastomer (World Precision Instruments Inc.) following removal of pipette. C57BL6/J mice were injected with *pAAV(PHP.eB)-BiPVe4-tdTom-tdTom_nls_* (Addgene 213936) in bilateral dorsal CA1 and CTX (100 nL at each site) at the following coordinates: -2.06 mm caudal and ± 1.60 mm lateral from bregma, 1.48 mm (CA1) and 0.81 mm (CTX) deep from the dura. PV^Cre^:Oprm1^fl/fl^ mice were injected bilaterally (70 nL at each site) with *pAAV(AAV9)-Ef1a-DIO-C1V1(t/t)-TS-mCherry* (Addgene 100061-AAV9, ≥ 1×10¹³ vg/mL) dorsal CA1 at the following coordinates: -2.06 mm caudal and ± 1.40 mm lateral from bregma, and 1.41 mm deep from the dura. Topical lidocaine, TAO, and ketoprofen were provided daily for 3 days following surgery. Mice were injected 2-4 weeks before experiments.

For perinatal injections, P0 mice were anesthetized by hypothermia. The scalp was wiped with 70% ethanol and a felt-tipped marker was used to mark the location of the injection over the lateral ventricles: 0.8 mm anterior to lambda and 0.8-1 mm lateral to the sagittal suture. Mice were bilaterally injected free-hand with 1 µL of solution containing *pAAV(AAV5)-pCAG-FLEX-EGFP-WPRE* (Addgene 51502-AAV5, ≥ 7 × 10^12^ vg/mL) and 0.05% trypan blue in the lateral ventricle, 2.5 mm below the surface of the scalp, using a 32G microliter Neuros Syringe (Hamilton). Mice were placed on a heating pad until movement returned and were rubbed with bedding material from the home cage before being returned to the mother.

### Viral Injections (macaque)

Intracranial macaque injections were targeted using stereotaxic coordinates derived from MRI and delivered using a needle guide for enhanced accuracy^110^. Surgeries were performed under aseptic conditions in a fully equipped operating suite. *pAAV(PHP.eB)-S5E2-ChR2-mCherry* (Addgene 135634-PHPeB, ≥ 1×10¹³ vg/mL) was injected into HPC and M1 of 6 rhesus macaques. Within HPC, 30-50 μL of total virus was injected, with 15-25 μL of virus injected at each of 2 locations spaced approximately 2 mm apart in the antero-posterior plane, caudal to the level of the uncus. Within M1, 30-40 μL of total virus was injected, with 10 μL of virus injected at each of 3-4 locations spaced approximately 2 mm apart, targeted via direct visualization.

For brain extraction, 6-8 weeks after virus injection, animals were sedated with ketamine/midazolam (ketamine 5-15 mg/kg, midazolam 0.05-0.3 mg/kg) and maintained on isoflurane. A deep level of anesthesia was verified by an absence of response to toe-pinch and absence of response to corneal reflex. Prior to brain removal and blocking, 5 macaques for electrophysiology were transcardially perfused with ice-cold sucrose-substituted artificial cerebrospinal fluid (aCSF) containing in mM: 90 Sucrose, 80 NaCl, 3.5 KCl, 1.25 NaH_2_PO_4_, 24 NaHCO_3_, 10 Glucose, 0.5 CaCl, 4.5 MgCl_2_ saturated with carbogen (95% O_2_, 5% CO_2_), with osmolarity 310-320 Osm. One macaque was fixed-perfused for IHC, euthanized following AVMA guidelines, and transcardially perfused with heparinized saline followed by a solution of 4% paraformaldehyde (PFA) in 0.1M phosphate buffer. The brain was removed and cryoprotected through an ascending series of glycerol solutions. The cryoprotected tissue was then frozen in isopentane and serially sectioned (at 40 µm) using a sledge microtome. Series with every tenth section (400 µm apart) of free-floating sections were processed for IHC.

### Morphine Injections

PV^Cre/+^:ChR2^fl/+^ mice were injected subcutaneously with morphine or saline in three treatment regimens, including a bolus injection (15 mg/kg), a chronic treatment with increasing doses over 6 days (15, 20, 25, 30, 40, 50 mg/kg), and a withdrawal group receiving the chronic treatment with delayed acute slice preparation. Morphine Sulfate (4 mg/mL, Hikma Pharmaceuticals, NDC:0641-6125) was diluted in saline so that each mouse received the same total volume (300 mL). Injections were performed at 9 AM each day, with acute slices prepared one hour after the final injection, except for the withdrawal group in which 72 hours passed. Mice were injected on a staggered alternating schedule so that 1 mouse was recorded from each day, alternating between saline and morphine.

### Acute Slice Preparation

Adult mice were anesthetized with isoflurane and rapidly decapitated. Brains were dissected, blocked, and sectioned in iced oxygenated sucrose-substituted aCSF. Horizontal or coronal slices (300 µm) were sectioned on a VT-1200S vibratome (Leica Microsystems), then transferred to a submerged incubation chamber containing oxygenated warmed (32-34 °C) sucrose-substituted aCSF for 30 minutes then maintained at room temperature for the duration of the day. Slices were allowed to recover post-sectioning for at least 1 hour before recording. Macaque and human samples were sectioned identically but maintained for up to 72 hours in room temperature oxygenated sucrose-substituted aCSF, with solution changes every 24 hours. Juvenile mice (<P21) were sectioned identically except high-Mg^2+^ instead of sucrose-substituted aCSF was used for dissection, blocking, sectioning, and incubation, containing in mM: 130 NaCl, 3.5 KCl, 1.25 NaH_2_PO_4_, 24 NaHCO_3_, 10 Glucose, 1 CaCl, 5 MgCl_2_, with osmolarity 300-310 Osm. Slices for GDP recordings were sectioned at 500 µm.

### Slice Electrophysiology

Slices were transferred to an upright microscope (Olympus BX51Wl) and perfused with oxygenated extracellular aCSF containing in mM: 130 NaCl, 3.5 KCl, 1.25 NaH_2_PO_4_, 24 NaHCO_3_, 10 Glucose, 2.5 CaCl, 1.5 MgCl_2_, with osmolarity 300-310 Osm, with a flow rate of 2-3 mL/min at a temperature of 30-33 °C. Individual cells were visualized with a 40x objective using fluorescence and IR-DIC microscopy. Electrodes were pulled from borosilicate glass (World Precision Instruments) to a resistance of 3-5 MΩ with a vertical pipette puller (Narishige PC-10). Whole-cell patch-clamp recordings were made with a Multiclamp 700B amplifier (Molecular Devices), with signals digitized at 20 kHz (Digidata 1440A, filtered at 3 kHz). Recordings were made using a Windows 10 computer with pClamp 10.7 (Molecular Devices). In voltage-clamp recordings, uncompensated access resistance (R_A_) was monitored consistently with 5 mV voltage steps. Any recordings in which R_A_ deviated by more than 20% were discarded, as were any with unstable leak current.

Direct response to opioids were recorded in fluorescently-identified cells with a standard K-Gluconate internal containing in mM: 150 K Gluconate, 0.5 EGTA, 3 MgCl_2_, 10 HEPES, 2 ATP•Mg, 0.3 GTP•Na_2_, and 0.3% biocytin (pH corrected to 7.2 with KOH, 285-290 Osm). Cells were held in voltage-clamp at - 50 mV, and 100 nM DAMGO (μOR selective agonist) and 500 nM CTAP (μOR selective neutral antagonist) applied. GPCR function was occluded with an alternate internal solution replacing the 0.3 mM GTP•Na_2_ with 1 mM GTPγS • Li_4_, allowing 15 min of baseline before applying DAMGO/CTAP.

Recordings of inhibitory light-evoked and spontaneous inhibitory post-synaptic currents (leIPSCs and sIPSCs) were made in visually-identified PCs held in voltage-clamp at -70 mV, with a high Cl^-^ K-Gluconate internal (calculated Cl^-^ reversal = -24 mV), containing in mM: 100 K Gluconate, 50 KCl, 0.5 EGTA, 3 MgCl_2_, 10 HEPES, 2 ATP•Mg, 0.3 GTP•Na_2_, 4 QX314, and 0.3% biocytin (pH corrected to 7.2 with KOH, 285-290 Osm). A brief (0.5-2 ms) 470 nm light pulse train (20x at 10 Hz) illuminated a small patch surrounding the recorded cell every 10 s. The light intensity was kept consistent between cells as much as possible, but as every recording was internally normalized to baseline, greater importance was placed on ensuring the leIPSC amplitude was detectable above noise and non-saturating (100-1000 pA). GABA_A_ currents were isolated by washing into the bath 10 µM DNQX, 50 µM DL-APV, and 1 µM CGP 55845, to eliminate the contributions of AMPA, NMDA, and GABA_B_ receptors, respectively. After a stable baseline was reached (5+ min), the opioids of interest were applied in the following nM concentrations: 100/1000 DAMGO, 500 CTAP, 500 DPDPE (δOR selective agonist), 100 naltrindole (δOR selective neutral antagonist), 10000 morphine (μOR/δOR agonist), and 1000 naloxone (μOR/δOR antagonist/inverse agonist). Each drug was applied for at least 5 min. Opioid concentrations were selected from prior publications^21,52,58,111^. Detailed dose-response experiments were not conducted.

Intrinsic membrane and firing properties were recorded in cells with a standard K-Gluconate internal. Resting membrane potential (RMP) was measured in a tight cell-attached configuration, voltage-clamped to +60 mV during 100 ms voltage ramps (from +100 to -200 mV) every 5 s and followed after break-in and a whole-cell current-clamp configuration with holding current I = 0. Membrane time constant (tau) was measured by 20 repeated 400 ms hyperpolarizing current pulses of -20 pA. Input resistance (R_in_) was determined by recording the response to 20 increasing current pulses (2 s duration from -50 pA with increasing 5 pA increments). Sag index was measured during 800-1000 ms current steps in which the peak voltage V_peak_ = -100 mV. Action potential (AP) and after-hyperpolarization (AHP) characteristics were measured during 800-1000 ms current steps near spike threshold. Mean max firing and spike adaption properties were measured during 800-1000 ms current steps at the maximum depolarizing current that preceded depolarization block. The range of current steps was dependent on input resistance and set for each cell to cover voltage responses from -100 mV to threshold to maximum firing.

Paired Tac1→PC recordings were conducted by targeting and characterizing intrinsic parameters of a fluorescently-identified Tac1 cell with a standard K-Gluconate internal, followed by targeting a non-labeled PC and recording in voltage-clamp at -70 mV with a high Cl^-^ K-Gluconate internal. A train of depolarizing current pulses (20x 2 ms 1-2 nA at 100 Hz) was injected into the pre-synaptic Tac1 cell to elicit unitary IPSCs (uIPSCs) in the post-synaptic PC. Multiple post-synaptic PCs were targeted within the same slice until detectable uIPSCs were observed, after which the drug of interest (100 nM DAMGO) was applied.

Giant depolarizing potential associated currents (GDP-Is) were recorded intracellularly, by recording visually-identified PCs voltage-clamped to 0 mV in standard K-Gluconate. Optogenetic inactivation of Tac1 cells was achieved by sustaining a light pulse for 2-3 min at 580 nm in ArchT-expressing mice.

### Electrophysiology Analysis

All electrophysiology analysis was conducted in Clampfit 10.7 (Molecular Devices) and processed in Microsoft Excel. Drug responses of the holding current were calculated as the final 2 min average for each drug condition. leIPSC amplitudes recorded in PCs were calculated as the final ten sweep average for each drug condition. sIPSCs were detected using a template search and averaged for the final 2 min of each drug condition. GDP-Is were detected using a threshold set above spontaneous sIPSCs and with a minimum event duration of 25 ms. GDP-Is were quantified for the final 2 min of each drug condition, and only recordings with a rate ≥ 2 events/min in the baseline period were included. GDP-Is recorded in S1 occurred at a significantly lower rate, and thus were conducted over longer recordings, quantified in the final 4 min of each drug condition and included if ≥ 1 event/min within the baseline period.

Intrinsic membrane and firing properties were analyzed as follows: RMP was calculated as the linearly-extrapolated intersection point along the voltage ramp as previously described^112^, or the mean voltage in I = 0 whole-cell recordings. Tau was determined by fitting the mean voltage response to hyperpolarizing current injections with an exponential function. R_in_ was determined by taking the slope of a linear regression of the change in voltage in response to increasing current pulses around RMP. The sag index was determined by measuring the maximum hyperpolarization voltage deflection (V_peak_) from baseline (V_rest_) at the onset of the current pulse, and the steady-state voltage deflection (V_ss_) in the last 200 ms of the current pulse. The sag index was calculated as (V_rest_ – V_ss_) / (V_rest_ – V_peak_). AP threshold was measured as the voltage at which the slope exceeds 10 mV/ms, and half-width as the width of the AP (ms) at half the maximum voltage amplitude. AHP amplitude and time was measured at the peak hyperpolarization voltage deflection from AP threshold. The maximum firing frequency was measured as the mean frequency during the maximum 800-1000 ms AP train, while the accommodation ration was measured as the ratio of the final (last 3 spikes) over the initial (first 3 spikes) instantaneous frequency.

### Single nucleus RNA sequencing (snRNAseq)

Detailed acquisition and analysis procedures of single nucleus RNA sequencing (snRNAseq) datasets of P10 and P28 hippocampal interneurons will be included in a subsequent publication. In brief, hippocampal tissue was dissected from Dlx5/6^Cre^:CAG-Sun1/sfGFP mice. Nuclei were isolated as previously described^113^ and subsequently sorted on a Sony SH800S cell sorter for GFP+ nuclei. snRNAseq libraries were prepared using the Chromium single cell 3’ library and gel beads kit (10x genomics, Cat# PN-1000075). CellRanger v7.0.0 (10x Genomics) was used with default parameters to map snRNAseq data to the mouse reference genome (mm10) provided by 10x Genomics. The snRNAseq data was then processed and annotated using the pipeline as previously described^114^.

### In situ Hybridization (ISH) RNAscope

To prepare snap frozen tissue for ISH, freshly dissected brains or brain blocks were submerged into 2-methylbutane that was pre-chilled on a dry ice/ethanol bath for 1 minute. Tissue was removed, wrapped in foil, and stored at -80 °C. 10 μm frozen sections were made using a Leica Cryostat, mounted on slides (ThermoFisher Scientific) and stored at -80 °C.

Target probes were designed and manufactured by Advanced Cell Diagnostic (ACD): Mm-Pvalb (Cat# 421931), Mm-Oprm1-C2 (Cat# 315841-C2), Mm-Oprd1 (Cat# 427371), Mmu-PVALB (Cat# 461691), Mmu-OPRM1-C2(Cat# 518941-C2). ISH was performed following the RNAscope Multiplex Fluorescent v2 Assay instructions provided by ACD. Briefly, frozen thin sections were post-fixed in 4% PFA and dehydrated sequentially in 50%, 70% and 100% ethanol. After H_2_O_2_ and Protease IV treatment, sections were incubated with probes at 40 °C for 2 hours. Probed signals were detected using RNAscope Multiplex Detection v2 kit (ACD, Cat# 323100). Opal 520 (AKOVA, Cat# OP-001001) and Opal 570 (AKOVA, Cat# OP-001003) were used to detect C1 and C2 probes, respectively. After DAPI staining, sections were covered using ProLong Gold Antifade Mountant (ThermoFisher Scientific, Cat# P10144) and cured in darkness before imaging.

### EdU Protocol

Timed pregnant C57BL/6 Mice (Charles River) were injected once each with 1.25 mg of EdU (Thermo Fisher, Cat# A10044) at E11, E12, E13, E14, and E15 via intraperitoneal injection. Upon reaching P30-32, mice were anesthetized with sodium pentobarbital (Euthasol) via intraperitoneal injection and transcardially perfused with 1x PBS followed by 4% PFA. Brains were then dissected and post-fixed in 4% PFA overnight at 4° C. PFA-fixed brain samples were cryopreserved in 30% (w/v) sucrose and sectioned into 20 µm coronal slices using a sliding microtome (Leica). Brain slices were preserved in Storage Buffer, comprising 28% (w/v) sucrose, 30% (v/v) ethylene glycol in 0.1 M sodium phosphate buffer, at -80 °C until further processing. mRNA transcripts were detected using the RNAscope Multiplex Detection v2 kit (ACD, Cat# 323100), following the manufacturer’s protocol. The RNAscope catalogue probes used included *Oprm1* (#315841-C2), *Oprd1* (#427371), and *Pvalb* (#421931-C3). Following RNAscope, EdU was detected using the Click-iT EdU imaging Kit (Thermo Fisher, Cat# C10340). For each time point analyzed, two animals from the same litter were used, and two sections were taken from each brain. Images of RNAscope ISH experiments were collected using an upright confocal microscope (Zeiss LSM800) with a 10x objective (Plan-Apochromat 10x/0.45 M27), with stitching performed using the Zeiss software.

### RiboTag

This assay was performed as previously described^44–46^. RNA bound with anti-HA immunoprecipitates and RNA from bulk tissue were purified using RNeasy Plus Micro Kit (Qiagen, Cat# 74034) and the quality of RNA was measured using RNA 6000 Pico kit (Agilent, Cat# 5067-1513) and 2100 Bioanalyzer system (Agilent, Cat# G2939BA). cDNA libraries were constructed from 250 pg RNA using the SMARTer Stranded Total RNA-Seq. Kit v2 (Takara Bio, Cat# 634411) from samples with RNA Integrity Numbers > 6. Sequencing of the libraries was performed on the Illumina HiSeq 2500, at 50 million 2 × 100 bp paired-end reads per sample. ∼75% of reads were uniquely mapped to genomic features in the reference genome. Bioconductor package DESeq2^109^ was used to identify differentially expressed genes (DEG). This package allows for statistical determination of DEGs using a negative binomial distribution model. The resulting values were then adjusted using the Benjamini and Hochberg method for controlling the false discovery rate.

### Immunohistochemistry (IHC)

For mice younger than P10, brain tissues were dissected and drop fixed in 4% PFA for 24 hours at 4 °C. Mice older than P10 were transcardially perfused using 4% PFA and dissected brain tissues were post-fixed in 4% PFA for 24 hours at 4 °C. Fixed brain tissues were thoroughly washed in 1x phosphate buffer (PB) followed by cryopreservation using 30% sucrose. 50 μm coronal sections were made on a frozen microtome. Brain slices were washed with 1x PB at room temperature for 1 hour with 2-3 changes of 1x PB. To perform floating section IHC, brain slices were blocked and permeabilized in Blocking Solution (1x PB + 10% goat serum + 0.5% Triton X-100) at room temperature for at least 2 hours. Primary antibodies were diluted using Antibody Solution (1x PB + 1% goat serum + 0.1% Triton X-100). Blocked brain slices were incubated in primary antibodies at 4 °C for 48 hours. After wash with 1x PB at room temperature for 15 minutes with 3 repeats, brain slices were incubated in secondary antibodies diluted with Antibody Solution at room temperature for 1 hour. After wash with 1x PB at room temperature for 15 minutes with 3 repeats, brain slices were mounted on gelatin coated slides followed by air drying, cover-slipped with Prolong Diamond Antifade mountant with DAPI (ThermoFisher Scientific Cat# P36962), cured in darkness, and imaged. Primary and secondary antibodies with product numbers listed in Key Resources Table, with working dilutions of 1:1000 for all except primary Rabbit anti-pIκBα (Cell Signaling Technology Cat# 2859, RRID:AB_561111) and secondary CF647 Donkey anti-Mouse (Biotium Cat# 20046-1, RRID:AB_10853778), both at 1:500.

### Morphological Reconstruction

Slices containing biocytin filled cells were drop fixed in 4% PFA overnight at 4 °C. Fixed slices were then washed in PBS, permeabilized with 0.3% Triton X-100 and incubated with Alexa-488 or Alexa-555 conjugated streptavidin overnight (Thermo Fisher Scientific, Cat# S11223, S21381). When additional staining was desired, tissue was incubated in Blocking Solution (Carrier Buffer + 10% goat serum + 0.5% Triton X-100) at room temperature for at least 2 hours. Primary antibodies were diluted using Carrier Buffer (1% bovine serum albumin + 1% goat serum in PBS) containing 0.5% Triton X-100 for 3 hours at room temperature. After washing with 1x PBS at room temperature for 15 minutes with 3 repeats, tissue was incubated with secondary antibodies diluted with Carrier Buffer and Triton at room temperature for 2 hours. After incubation, slices were incubated in 1 μg/ml DAPI (Millipore Sigma, Cat# D9542), then underwent multiple washes, cryopreserving in 30% sucrose and were resectioned to 70 μm using a freezing microtome (Microm, Whaltham, MA). Slices were then mounted on glass slides (Fisher Scientific, Fisherbrand Superfrost Plus) using Mowiol as a mounting medium. Confocal images of labelled cells were attained on Zeiss LSM 710 and LSM 900 microscopes using a 20x objective. Primary and secondary antibodies with product numbers listed in Key Resources Table, with working dilutions of 1:1000.

### IHC and ISH Image Acquisition and Analysis

IHC Images were captured on a Zeiss LSM 900 (confocal 20x and Airyscan 63x) and an Olympus VS200 slide scanner. Quantified images were captured from 2-5 sections for each subject. Microscope settings were kept consistent for all sections from each subject. Lower resolution confocal (20x) images used a broad z-stack (5 slices every 1.1 µm) which were maximum projected before subsequent analysis. High resolution Airyscan (63x) images used a detailed z-stack (60-70 slices every 0.15 µm). RNAscope images were first acquired on a SLIDEVIEW VS200 slide scanner (Olympus) mounted with an ORCA-Fusion Digital Camera (HAMAMATSU). Hippocampal and neocortical regions were scanned at 20x, guiding subsequent 63x confocal z-stack images at regions of interest (ROIs).

For each image, polygonal ROIs were drawn to demarcate hippocampal/neocortical layers and subregions. Mean intensity and correlation across channels for each ROI were then computed by a custom ImageJ (FIJI) macro (see Data and Code Availability). Cells were manually counted for colocalization analysis with the built-in FIJI multiselect tool independently for each channel in 3-4 sections for each subject. The above macro was used to subdivide the counts between ROIs, compute the average cell intensity, ROI intensity, and ROI area. The average signal-to-noise ratio (SNR), defined here as the mean cell intensity divided by the mean ROI SD, was determined to be significantly greater than 1 for all manual counts (see Table S1).

## QUANTIFICATION AND STATISTICAL ANALYSIS

Statistical analysis was conducted in Graphpad Prism 10. All data were tested for normality and lognormality with Shapiro-Wilk tests. Parametric tests were selected if all groups were normal. If at least one group was non-normal and all groups were log-normal, data were log-transformed prior to parametric testing. If data were neither normal nor log-normal, non-parametric tests were used whenever available. Throughout, summary data are presented as mean ± SEM with symbols representing individual values. Additional descriptive statistics and details of hypotheses testing are available for each figure in Table S1.

Cellular responses to drug treatments were analyzed with a 1-way repeated-measures design (ANOVA/Friedman, with Tukey’s/Dunn’s *post hoc* multiple comparisons). All statistical tests were conducted on raw non-normalized data, including plots showing responses normalized to baseline. Experiments missing at most one condition (CTAP/naltrindole) were included and analyzed with a mixed-effects model. Comparisons of the normalized DAMGO responses between regions were performed with a 1-sample t-test/Wilcoxon, comparing to 1 (baseline) and correcting sigma for the number of comparisons. Comparisons between different brain regions’ drug responses over time were analyzed with a 2-way design (region × time), with Šídák’s multiple comparisons restricted to comparisons between time-equivalent bins. In general, *post hoc* tests for 2-way analyses were restricted to comparisons of *a priori* interest, employing Šídák’s multiple comparisons tests.

## SUPPLEMENTAL INFORMATION

**Fig. S1:**
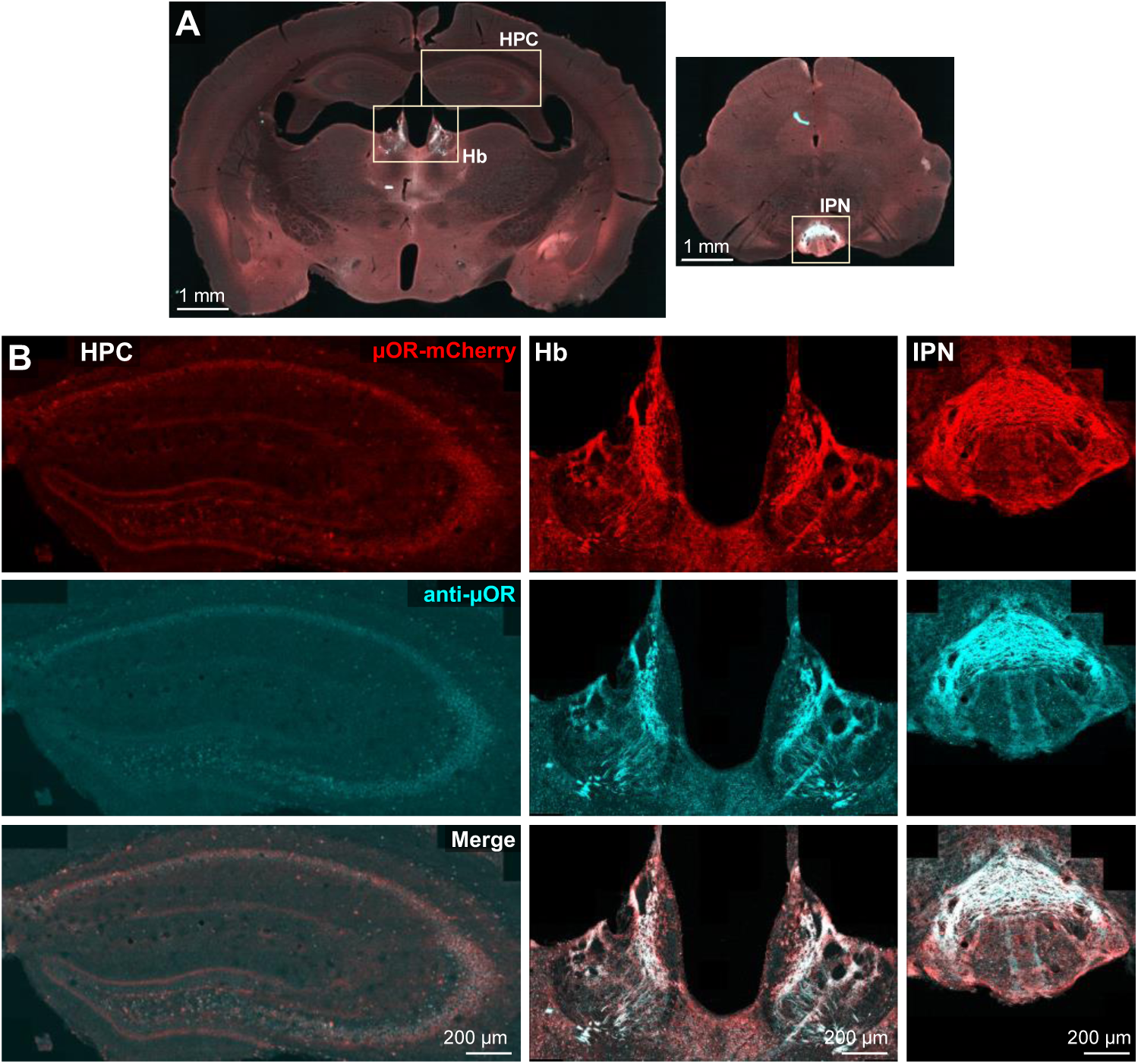
μOR-mCherry mouse model validation. (**A**) Coronal sections of IHC stain with antibody against µOR in µOR-mCherry mice (boosted with anti-RFP), bregma = -1.3 (*left*) and -3.3 (*right*). (**B**) Areas of high µOR expression at 20x magnification including hippocampus (HPC, *left*), habenula (Hb, *middle*), and interpeduncular nucleus (IPN, *right*). µOR antibodies (cyan) exhibited weaker somatic labeling than mCherry expression (red), but both channels appeared well colocalized. IHC performed in 2F P70 with consistent observations.

**Fig. S2:**
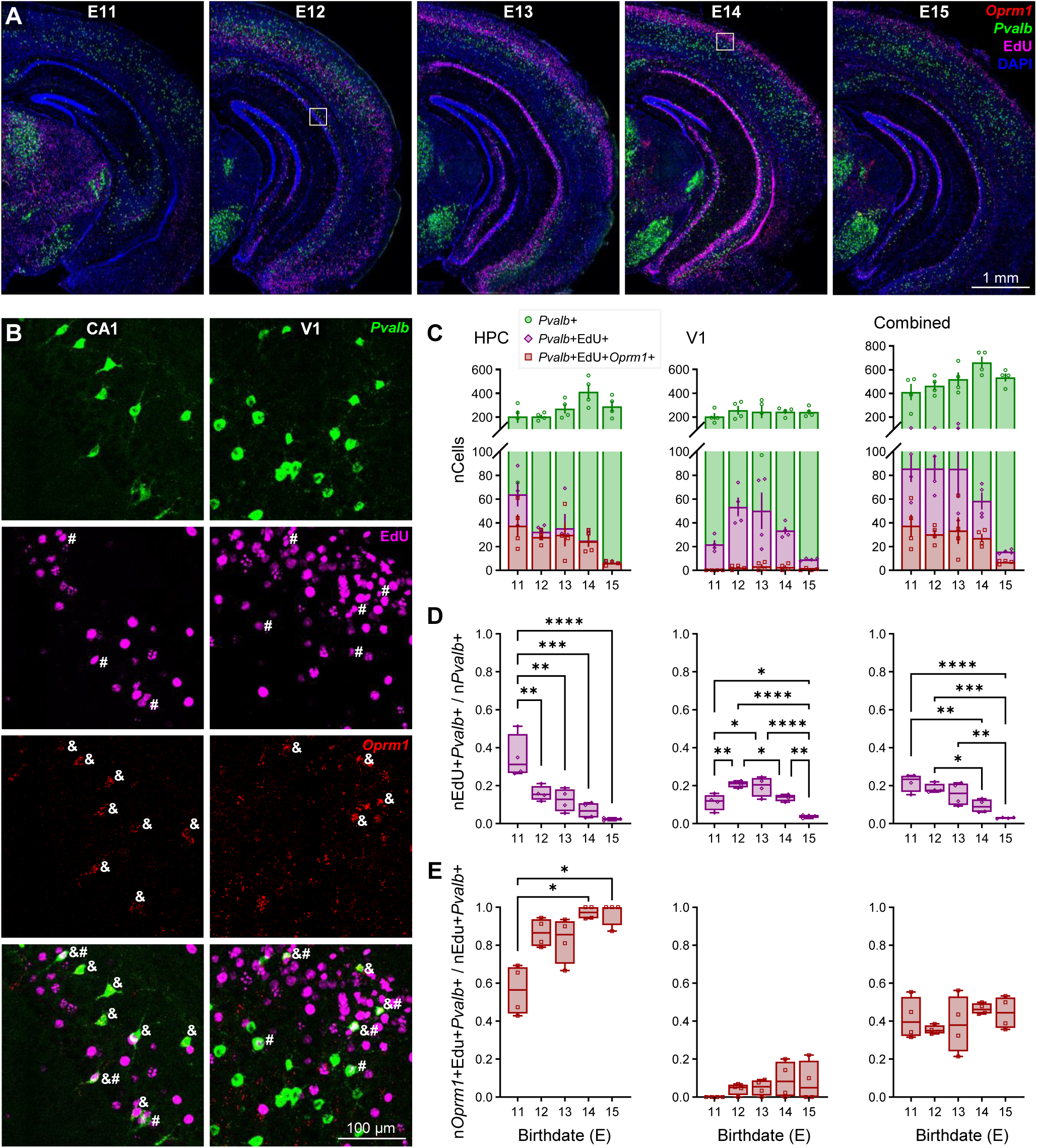
Earlier-born PV-INs are not more likely to express Oprm1. RNAscope for *Pvalb* and *Oprm1* in a total of 10 P28 mice, 2 each injected with EdU at E11, E12, E13, E14, and E15. (**A**) Representative coronal section with *Oprm1* (*red*), *Pvalb* (*green*), EdU (*magenta*), and DAPI (*blue*). Highlighted sections in CA1 (E12) and V1 L2/3 (E14) expanded in (**B**), with “#” markers signifying *Pvalb*+EdU+ co-labeled cells, “&” markers signifying *Pvalb*+*Oprm1*+ co-labeled cells, and “&#” markers signifying triple labeled *Pvalb*+Edu+*Oprm1*+ cells. (**C-E**) Triple colocalization quantification across embryonic birthdate E11-15 for HPC (*left*), V1 (*middle*) and CA1+V1 combined (*right*). Each animal included 2 technical replicates, with each quantified across left and right hemispheres. (**C**) Total number of counted *Pvalb*+, *Pvalb*+Edu+, and *Pvalb*+Edu+*Oprm1*+ cells. (**D**) Fraction of *Pvalb*+ cells co-labeled with EdU. (**E**) Fraction of *Pvalb*+EdU+ cells co-labeled with *Oprm1*. Asterisks represent Tukey’s *post hoc* comparisons after a significant effect of birthdate was found via 1-way ANOVA.

**Fig. S3:**
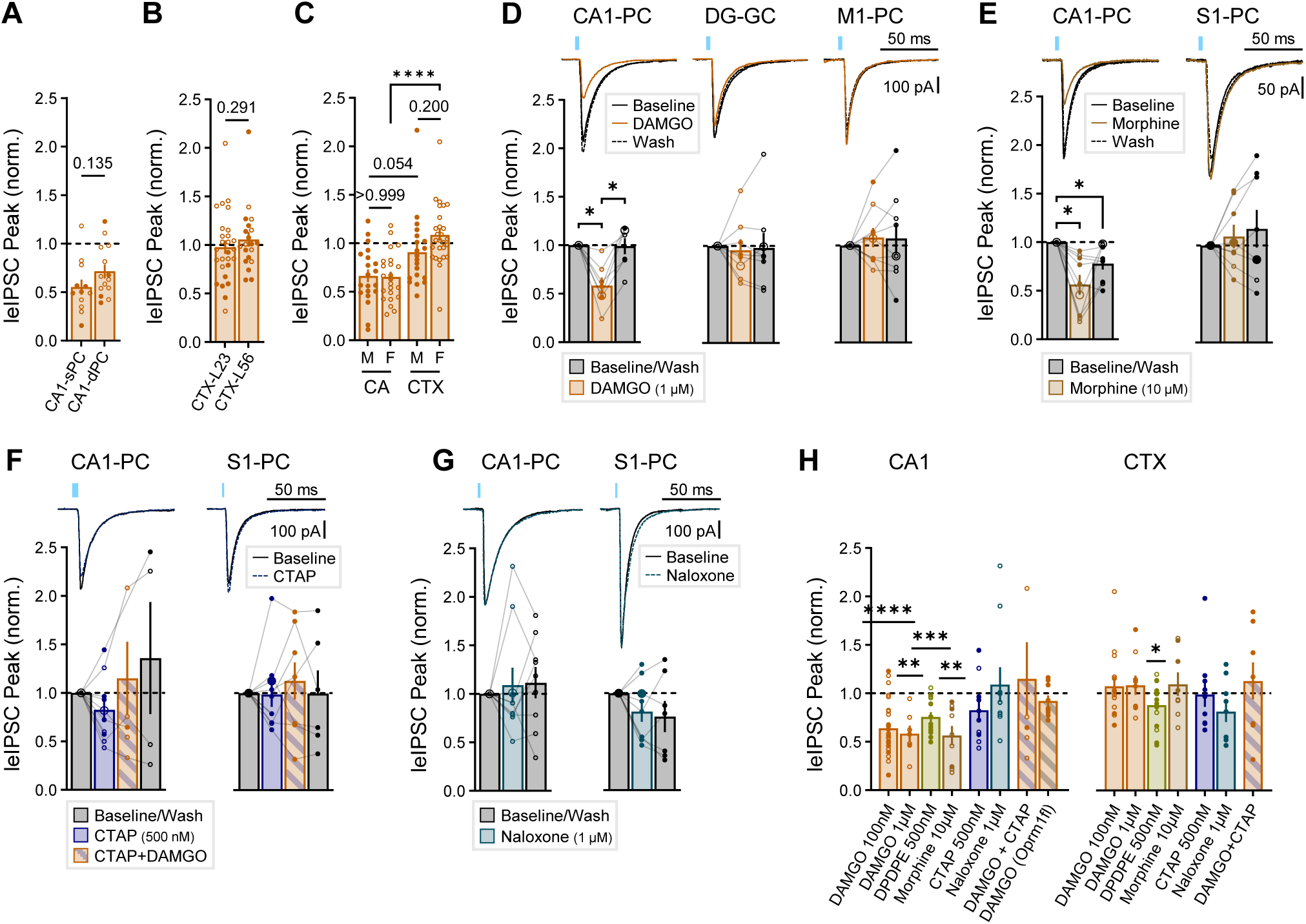
Additional PV-IN optogenetic experiments. Normalized leIPSC responses to bath administration of 100 nM DAMGO compared between (**A**) n_cell_ = 12 superficial and 14 deep CA1-PCs, (**B**) n_cell_ = 27 supragranular (L2/3) and 30 infragranular (L5/6) neocortical PCs, and (**C**) sex effects across n_cell_ = 20 (male), 23 (female) CA-PCs and 20 (male), 27 (female) CTX-PCs. Non-significant p-values and asterisks represent results of unpaired t-tests (*A*, *B*) and Šídák’s *post hoc* comparisons (*C*) after a significant effect of region was observed via 2-way ANOVA. (**D**-**G**) Mouse leIPSC experiments with (*top*) averaged example traces and (*bottom*) summary data for varied drug conditions. (**D**) DAMGO 1 µM: n_cell_ = 7 (CA1), 8 (DG), and 9 (M1) from n_mice_ = 5 (4F), age = P82-101 (P91 ± 4). (**E**) Morphine 10 µM: n_cell_ = 9 (CA1) and 8 (S1) from n_mice_ = 3 (2F), age = P33-118 (P83 ± 26). (**F**) CTAP 500 nM: n_cell_ = 10 (CA1) and 10 (S1) from n_mice_ = 2 (1F), age = P54, 60. (**G**) Naloxone 1 µM: n_cell_ = 10 (CA1) and 9 (S1) from n_mice_ = 3 (2F), age = P55-60 (P57 ± 2). Asterisks represent Tukey’s *post hoc* comparisons after a significant effect of treatment was found via 1-way repeated measure ANOVA/mixed model. (**H**) Baseline-normalized DAMGO response of PV-IN leIPSCs across all drug conditions in CA1 (*left*) and CTX (*right*), combining data from Fig. 3H*,I,N,U*, (*D-G*), and Fig. 4G. CTX represents S1 for all conditions except DAMGO 1 µM (M1) and DPDPE (V1). Asterisks represent significant deviations from the normalized baseline via 1-sample t-test.

**Fig. S4:**
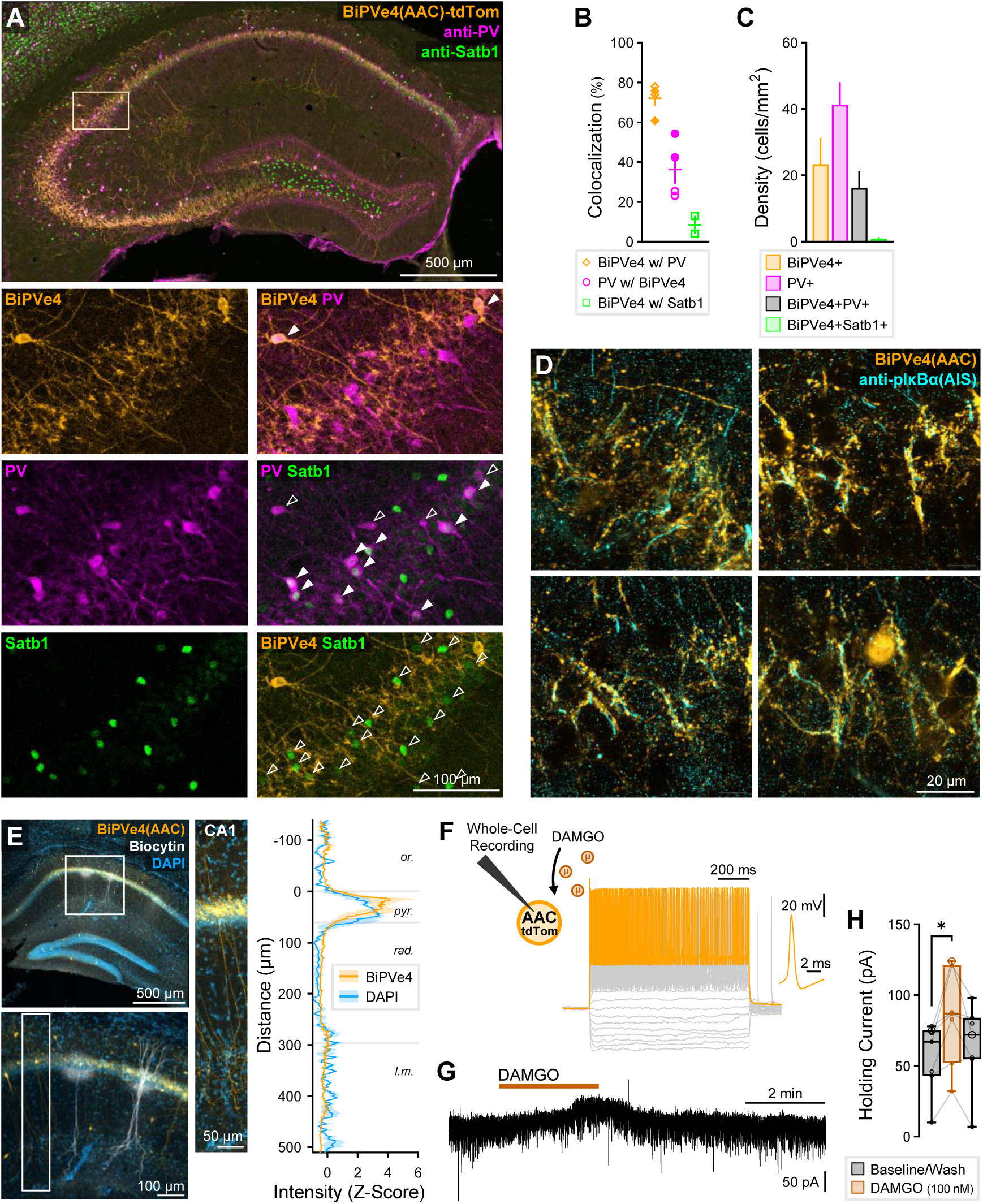
Validation of axo-axonic cell (AAC) specific BiPVe4 virus in rodent hippocampus. (**A**) IHC stain with antibodies against PV (*magenta*) and Satb1 (*green*) in 1M P155 C57BL6/J mouse injected retro-orbitally with BiPVe4-tdTom (*orange*, boosted with anti-RFP). In bottom zoomed images of CA2 region, open arrows indicate single marker labeling and closed arrows co-expression by indicated markers: (*from top to bottom*) BiPVe4+ & BiPVe4+PV+, PV+ & PV+Satb1+, Satb1+ & BiPVe4+Satb1+. (**B**) Colocalization quantification of triple IHC and (**C**) cell densities. Counts performed in 2 hippocampal sections each from two mice: 1M P155 C57BL6/J injected retro-orbitally with *BiPVe4-ChR2* and 1F P69 injected intra-cranially with *BiPVe4-tdTom*. (**D**) IHC stain with antibody against anti-pIκBα, labeling the axon initial segment (AIS, *cyan*) in 1F P44 injected retro-orbitally with *BiPVe4-tdTom*, showing close proximity of signals as expected from AACs. (**E**) Hippocampal section from same mouse as (*D*) after biocytin-labeling of recorded AAC (*white*). (*Right*) Intensity profile of 3 sections across CA1 layers, showing higher expression of BiPVe4 towards *oriens* side of *str. pyr.* (**F**) Schematic of whole-cell recordings of AACs illustrating membrane response to hyperpolaring/depolarizing current steps to measure intrinsic parameters (Table S2) and response to DAMGO. (**G**) Example trace showing change in holding current from DAMGO administration (rapid inward/outward currents reflect sEPSCs/sIPSCs). (**H**) Holding current summary data across baseline, DAMGO, and wash conditions for n_cell_ = 7 from 1F P44 injected intra-cranially with BiPVe4-tdTom. Asterisk represents Tukey’s *post hoc* comparison after a significant effect of treatment was found via 1-way repeated measure ANOVA.

**Fig. S5:**
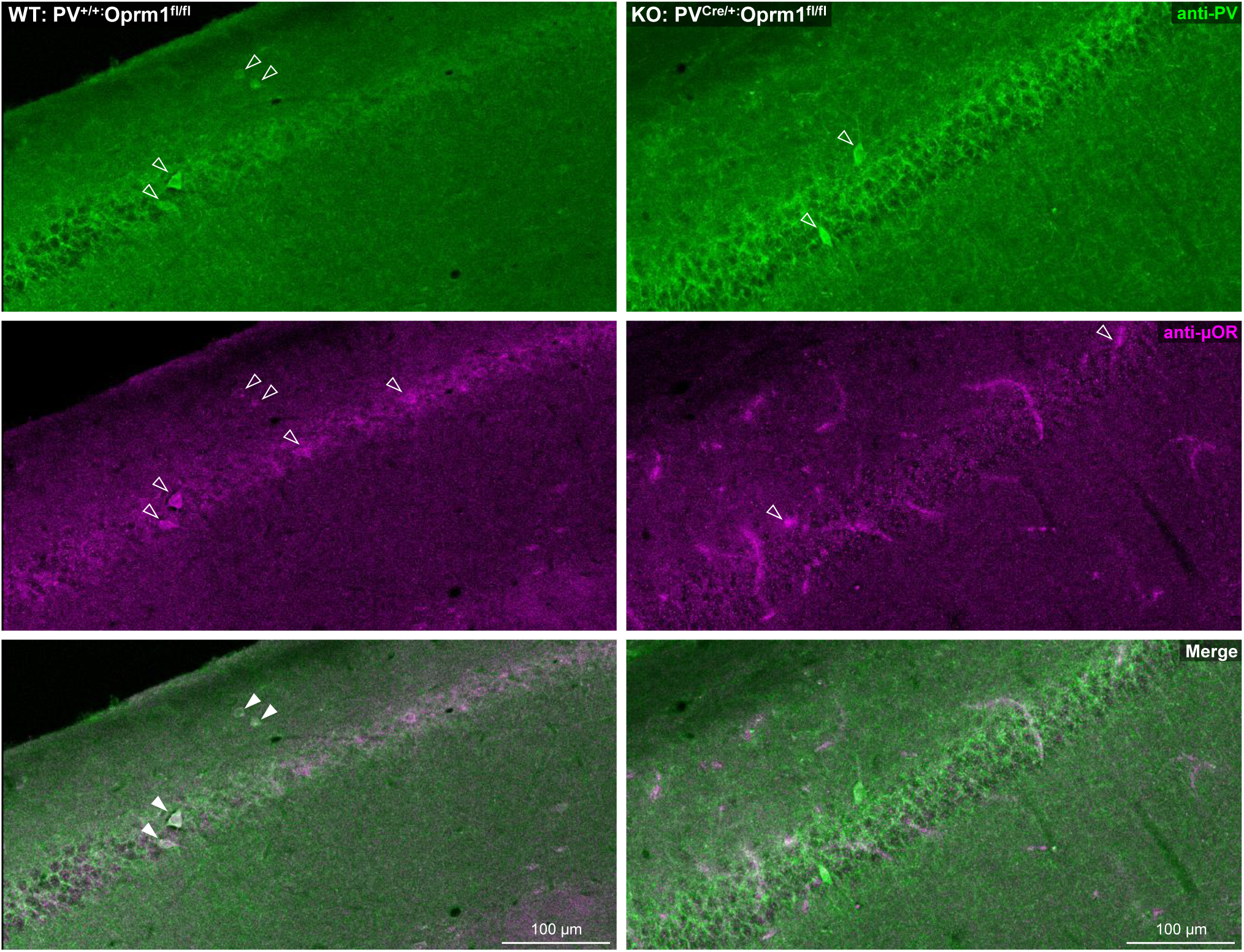
Validation of PV-IN selective Oprm1 knockout. Double IHC with antibodies against PV (*green, top*) and μOR (*magenta, middle*) in triple transgenic mice: PV-IN selective knockout (KO) mice (PV^Cre/+^:Oprm1^fl/fl^, 1M P77) and wild-type (WT) littermate controls lacking Cre-recombinase (PV^+/+^:Oprm1^fl/fl^, 1F P77). In the CA1 region pictured, the selective PV-IN KO (*right*) exhibits reduced μOR expression in identified PV-INs relative to littermate control (*left*). Open arrows indicate identified somata for respective signal, filled arrows indicate colocalized somata.

**Fig. S6:**
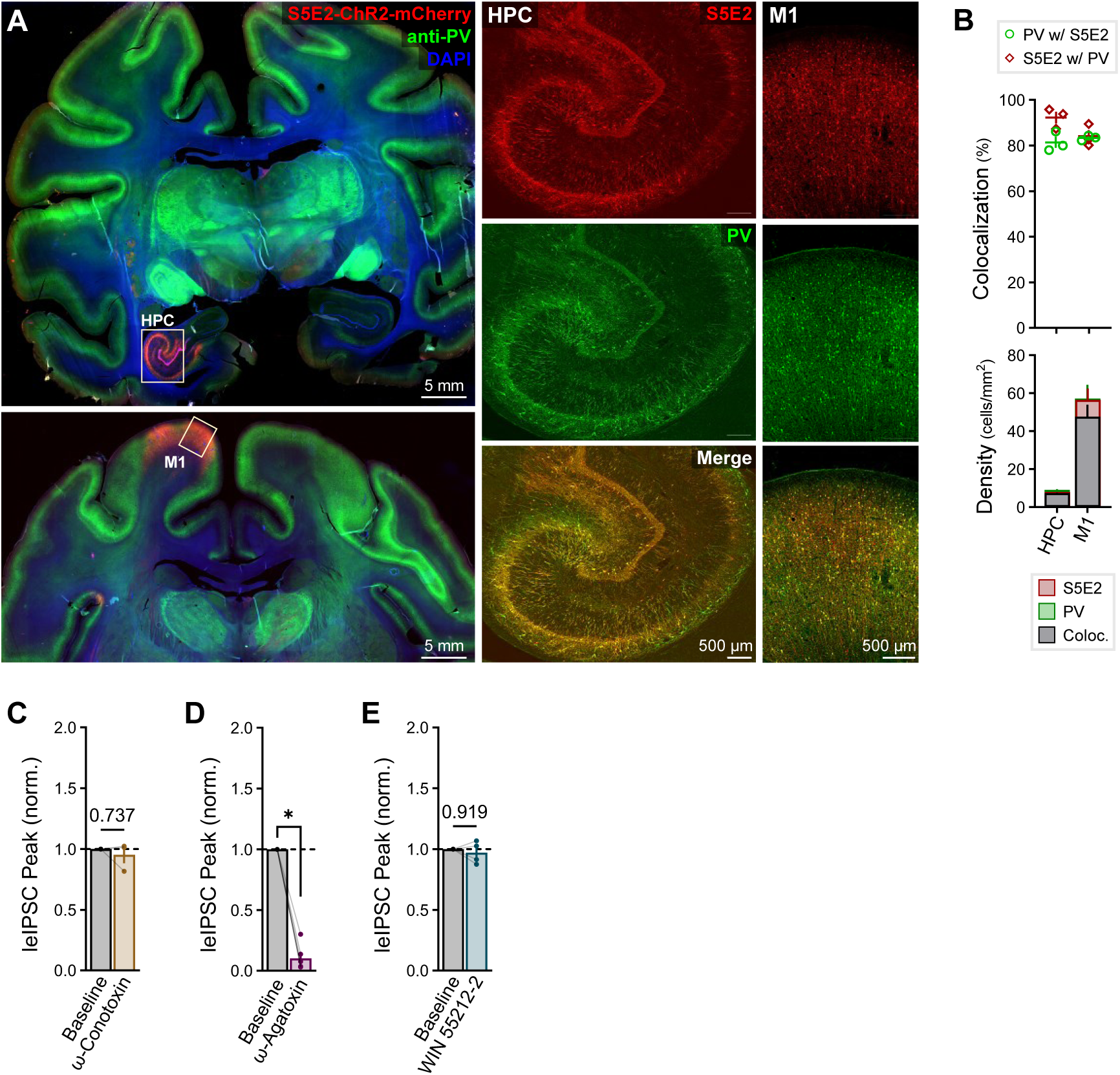
Validation of PV-specific S5E2 virus in rhesus macaques. (**A**) Whole-brain coronal slices of rhesus macaque injected with *pAAV(PHP.eB)-S5E2-ChR2-mCherry* (*red*) in HPC and M1, and stained for PV (*green*), to conduct colocalization analysis (*right*). (**B**) S5E2-PV colocalization quantified in 3 sections from 1F macaque, age = 11.7 years, quantified in 3 sections within each region, with (*top*) percentage of PV cells co-expressing S5E2 and vice versa, and (*bottom*) cell densities. (**C-D**) Normalized leIPSC responses in 1M macaque, age = 15.9 years, to bath administration of (**C**) 500 nM of the N-type Ca^2+^ channel blocker ω-conotoxin in n_cell_ = 3 CA-PCs (presynaptic channels used by CCK-INs^115,116^), (**D**) 250 nM of the P/Q-type Ca^2+^ channel blocker ω-agatoxin in n_cell_ = 6 CA-PCs (presynaptic channels used by PV-INs^116^), and (**E**) 5 μM of the synthetic cannabinoid agonist WIN 55212-2 in n_cell_ =4 CA-PCs (suppresses CCK-IN synaptic release through depolarization-induced suppression of inhibition (DSI)^117^). Non-significant p-values and asterisk represent results of paired t-tests.

**Fig. S7:**
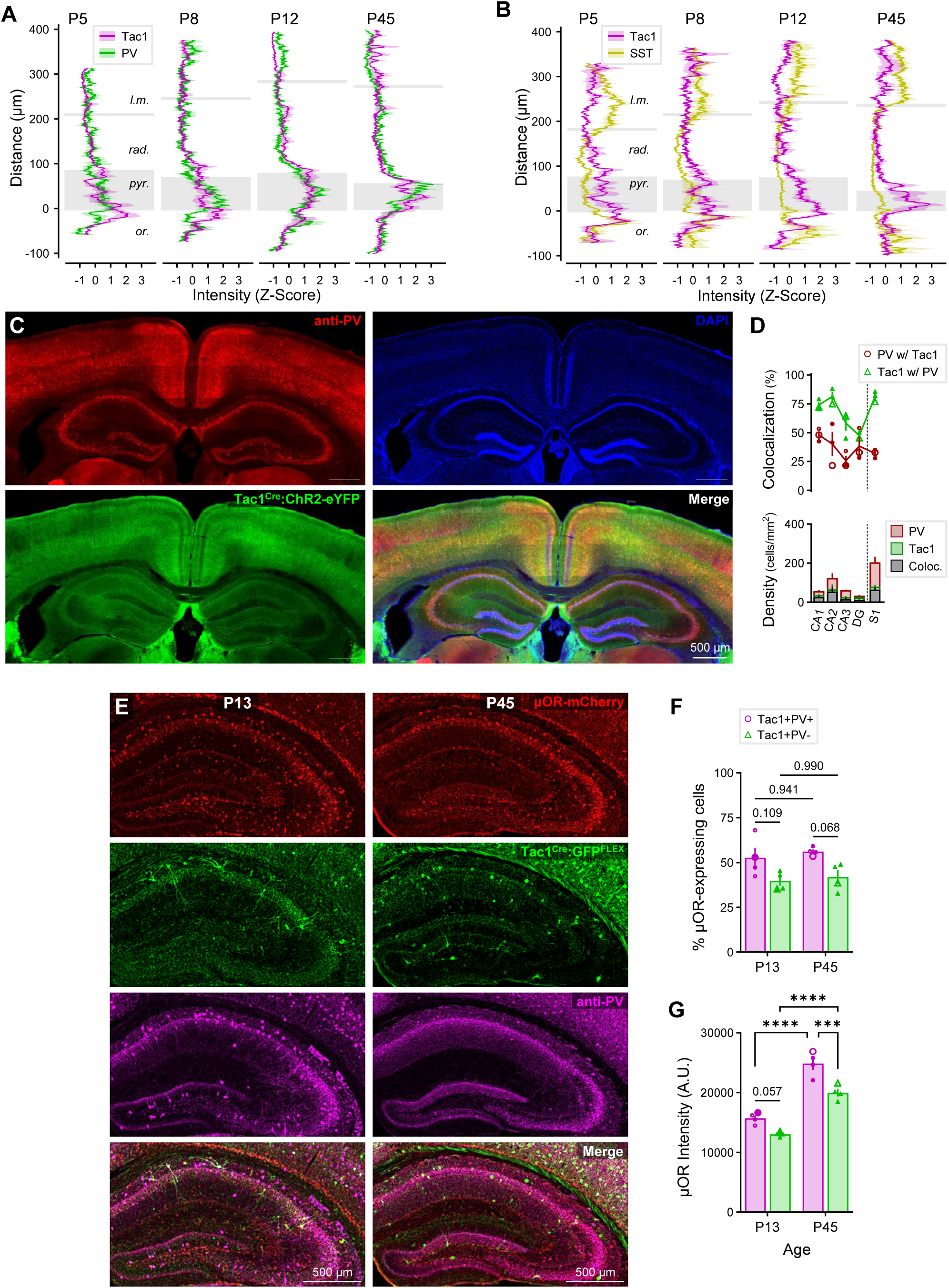
Additional IHC validation of Tac1^Cre^ mice as a model for immature PV-INs. (**A-B**) Quantified intensity profiles across hippocampal CA1 layers *l.m.*, *rad.*, *pyr.* and *or.* in P5-45 Tac1^Cre/+^:tdTom^fl/+^ mice, with distance = 0 set to *or.*-*pyr*. border. Intensity values for each section normalized by z-score prior to group averaging. (**A**) PV-Tac1 intensity profile average of n = 6 sections from n = 3 mice at each age (same data as example in Fig. 6D). (**B**) SST-Tac1 intensity profile average of n = 4 sections from n = 2 mice at each age (same data as example in Fig. 6F). (**C**) IHC stain for PV (*red*) in Tac1^Cre/+^:ChR2^fl/+^ (*green*), performed in 3 (2F) P37 mice, and (**D**) quantified in 3 sections for each mouse across HPC and S1, for (*top*) percent colocalization and (*bottom*) cell density. (**E**) IHC stain for PV in Tac1^Cre/+^:μOR-mCherry mice injected perinatally with *pAAV(AAV5)-pCAG-FLEX-EGFP-WPRE*, and thus triple-labeled for μOR (*red*), Tac1 (*green*), and PV (*magenta*). Stain replicated in 4 (2F) P13 juveniles and 4 (2F) P45 adults, and (**F-G**) quantified in 4 sections from each mouse, delineating Tac1+PV+ (*magenta*) and Tac1+PV-(*green*) populations (**F**) percentage of μOR-expressing cells and (**G**) μOR intensity (A.U. = arbitrary units). Asterisks represent Šidák’s *post hoc* comparisons after a significant effect of cell type was found via 2-way ANOVA.

**Fig. S8:**
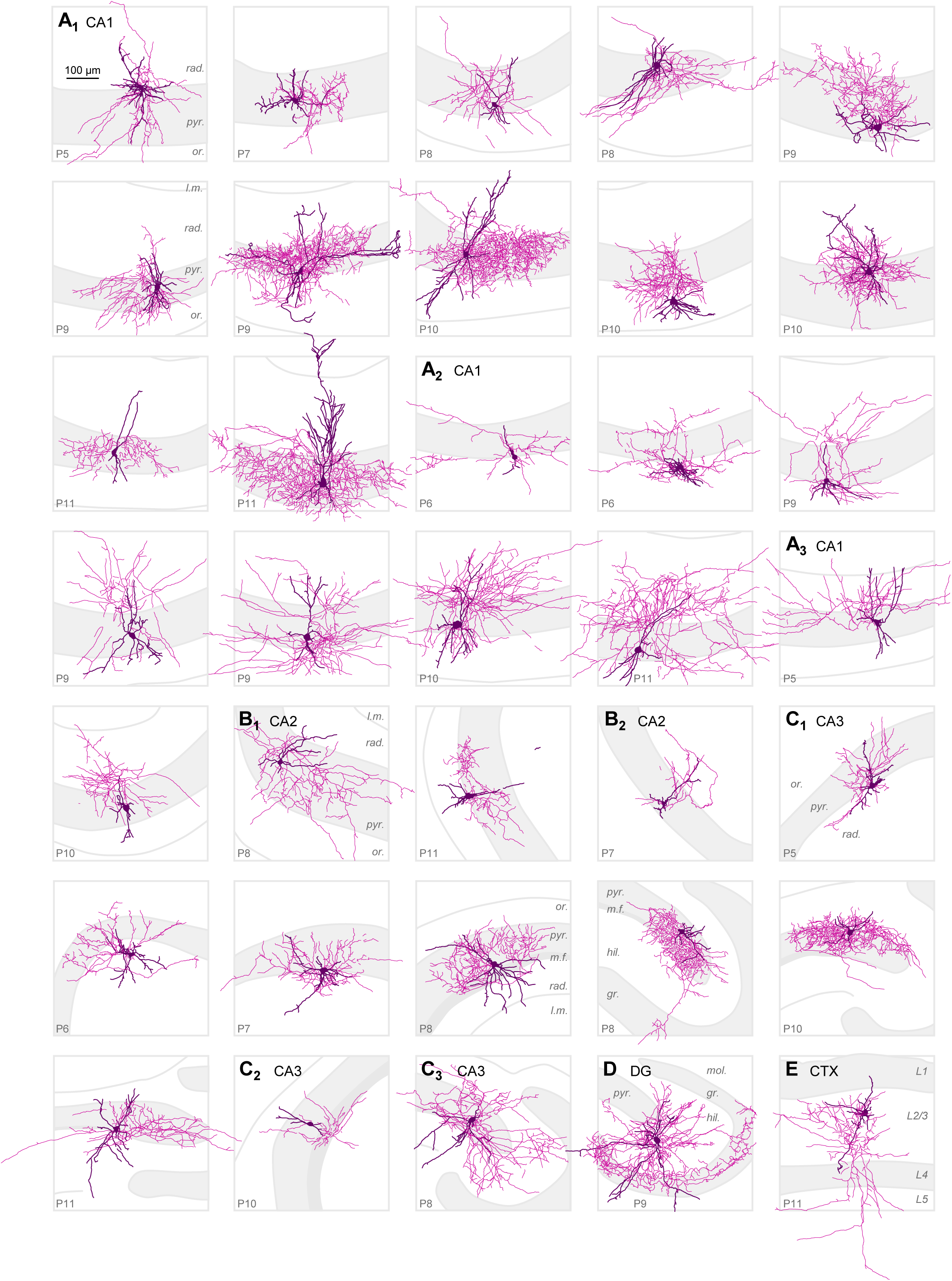
Morphological reconstructions of Tac1-INs in early development display perisomatic phenotypes consistent with immature PV-INs. Example morphological reconstructions of recorded Tac1 cells for (**A**) n_cell_ = 21 (CA1), (**B**) 3 (CA2), (**C**) 9 (CA3), (**D**) 1 (DG), and (**E**) 1 (CTX), recorded in 7 (P5-11) Tac1^Cre/+^:tdTom^fl/+^ mice (age indicated for each in lower left). Darker processes designate dendrites and soma, while lighter processes designate axons. CA1-3 layers: *stratum oriens* (*or*.), *pyramidale* (*pyr*.), *mossy fibers (m.f.)*, *radiatum* (*rad*.), *lacunosum-moleculare* (*l.m.*), DG layers: *molecular* (*mol*.), *granular* (*gr*.), *hilus* (*hil*.). Cells grouped by principal axonal target: primarily targeting *str. pyr.* and *gr.* consistent with BCs (**A_1_**, **B_1_**, **C_1_**, **D**), targeting both *str. or.* and *rad.* consistent with BSCs (**A_2_**, **B_2_**, **C_2_**), and primarily targeting superficial dendritic layers *rad.*, *l.m.*, and/or *hil.* (**A_3_**, **C_3_**). All reconstructions on same scale.

**Fig. S9:**
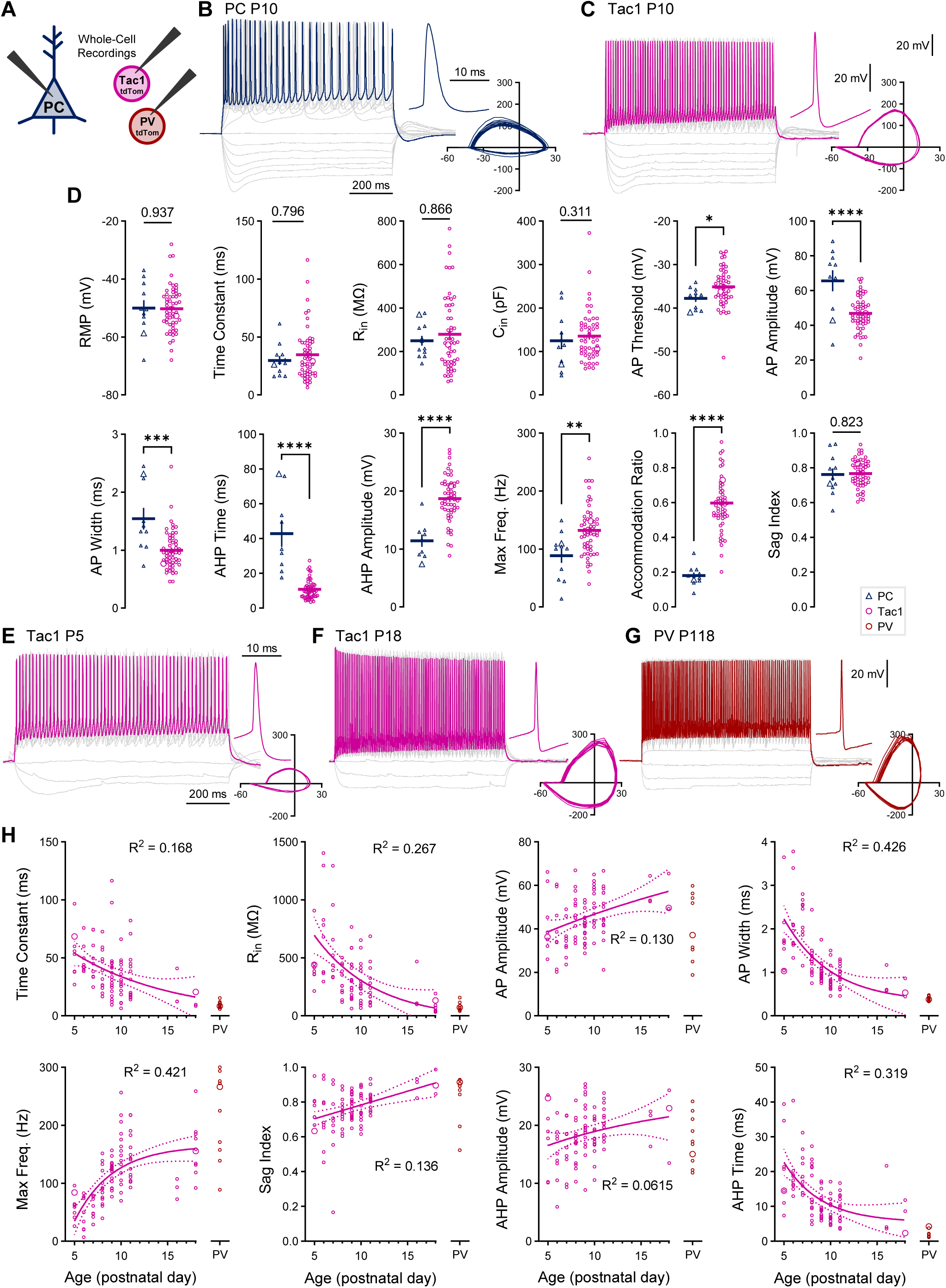
Intrinsic electrophysiological analysis of Tac1-INs reveals a fast-spiking phenotype consistent with immature PV-INs and distinct from PCs. (**A**) Schematic of whole-cell current clamp recordings of PCs/Tac1-INs from Tac1^Cre/+^:tdTom^fl/+^ mice and PV-INs from PV-tdTom mice to characterize intrinsic parameters. (**B-C**) Representative firing of (**B**) P10 PC and (**C**) P10 Tac1 cell, with gray traces showing response to range of hyperpolarizing and depolarizing current steps, and colored lines indicating maximum firing. Most hyperpolarized step = current required to take cell to -100 mV (*B*: -150 pA, *C*: -300 pA). Most depolarized step = current to take cell to maximum firing prior to depolarization block (*B*: +175 pA, *C*: +400 pA). (*Upper inset*) Single action potential (AP) on expanded time scale. All traces between cell types displayed on same scales. (*Lower inset*) dV/dt vs. V plot of ten successive APs to visualize AP shape differences. (**D**) Summary data for intrinsic parameters of hippocampal n_cell_ = 14 PC and 57 Tac1 cells from n_mice_ = 4 (P8-11) Tac1^Cre/+^:tdTom^fl/+^ mice. RMP = resting membrane potential, R_in_ = input resistance, C_in_ = input capacitance. Asterisks and non-significant p-values represent results of unpaired t-test/Mann Whitney. (**E-G**) Representative firing of (**E**) P5 Tac1, (**F**) P18 Tac1 cell, and (**G**) P118 PV cell (recorded in PV-tdTom mouse), with gray traces showing response to range of hyperpolarizing and depolarizing current steps, and colored lines indicating maximum firing. Most hyperpolarized step = current required to take cell to -80 mV (*E*: -50 pA, *F*: -100 pA, *G*: -350 pA). Most depolarized step = current to take cell to maximum firing prior to depolarization block (*E*: +175 pA, *F*: +350 pA, *G*: +550 pA). Same scales as (*B*, *C*). (**H**) Changes in select intrinsic parameters over time for both HPC and CTX Tac1 cells, plotting n_cell_ = 8 (1 P5), 9 (1 P6), 9 (1 P7),14 (1 P8), 20 (1 P9), 16 (1 P10), 14 (1 P11), 3 (1M P16), 10 (1M P18), and 10 PV (1F P118, 1F P392). Linear and exponential regressions performed on Tac1 data, with R^2^ values displayed, with PV data plotted for comparison.

**Fig. S10:**
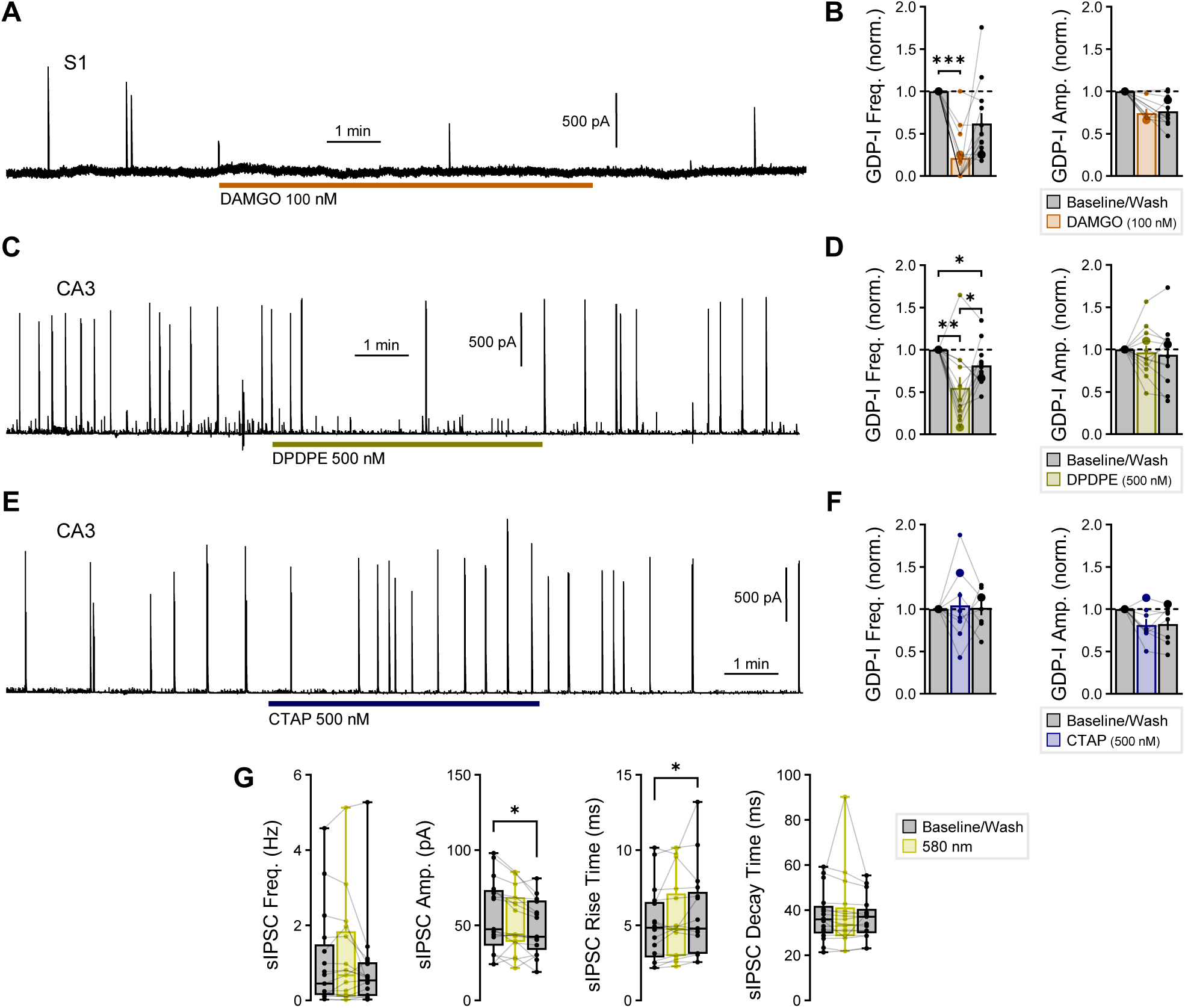
Additional GDP experiments. (**A**,**C**,**E**) Example traces of GDP associated currents (GDP-Is) recorded intracellularly and voltage-clamped to 0 mV in WT mice, with (**A**) 100 nM DAMGO applied for 10 min. in S1, (**C**) 500 nM DPDPE applied for 5 min. in CA3, and (**E**) 500 nM CTAP applied for 5 min. in CA3. (**B**,**D**,**F**) Summary data for GDP-I event frequency and amplitude for (**B**) n_cell_ = 13 S1-PCs from n_mice_ = 4 (P6-8), (**D**) n_cell_ = 12 CA3-PCs from n_mice_ = 3 (P6-7), and (**F**) n_cell_ = 8 CA3-PCs from n_mice_ = 4 (P7-8). (**G**) Summary data for spontaneous inhibitory currents (sIPSCs) detected between GDP-I events in n_cell_ = 17 from 6 Tac1^Cre/+^:ArchT^fl/+^ P5-8 mice (data in Fig. 7D*-F*), quantifying (*left to right*) sIPSC frequency, amplitude, rise time, and decay time. Asterisks represent Tukey’s/Dunn’s *post hoc* comparisons after a significant effect of treatment was found via 1-way repeated measure ANOVA /Friedman’s test.

***Table S1: Statistical details.***

Statistical details including summary statistics, normality tests, hypothesis testing, and *post hoc* comparisons, detailed in individual worksheets for each figure and supplemental figure.

***Table S2: Intrinsic Parameters.***

Intrinsic electrophysiological parameters recorded from 14 PCs (P8-11), 101 Tac1 cells (P5-18, delineated by subregion), 10 PV-INs (P118-392), and 14 AACs (P44).

## Notes

### Competing Interest Statement

The authors have declared no competing interest.

### Summary of Updates

Minor revisions. Additional recordings from younger P5-7 Tac1-tdTom mice, resulting in the electrophysiological characterization of 26 additional cells, 15 of which we morphologically reconstructed. Increased the number of biological replicates for PV-Tac1 and SST-Tac1 IHC experiments. Fig. 6, Fig. S6-9 saw minor revisions, with minor text revisions throughout.

